# The Respiratory Syncytial Virus M2-2 protein is targeted for proteasome degradation and inhibits translation and stress granules assembly

**DOI:** 10.1101/2023.01.02.522538

**Authors:** Orlando Bonito Scudero, Verônica Feijoli Santiago, Giuseppe Palmisano, Fernando Moreira Simabuco, Armando Morais Ventura

**Author notes:** Corresponding authors (OBS), (AMV).

## Abstract

The M2-2 protein from the respiratory syncytial virus (RSV) is a 10 kDa protein expressed by the second ORF of the viral gene M2. During infection, M2-2 has been described as the polymerase cofactor responsible for promoting genome replication. This function was first inferred by infection with a mutant virus lacking the M2-2 ORF, in which viral genome presented delayed accumulation in comparison to wild-type virus. In accordance with this phenotype, it has been recently shown that M2-2 promotes changes in interactions between the polymerase and other viral proteins at early stages of infection. Despite its well-explored role in the regulation of the polymerase activity, little has been made to investigate the relationship of M2-2 with cellular proteins. In fact, a previous report showed poor recruitment of M2-2 to viral structures, with the protein being mainly localized to the nucleus and cytoplasmic granules. To unravel which other functions M2-2 exerts during infection, we expressed the protein in HEK293T cells and performed proteomic analysis of co-immunoprecipitated partners, identifying enrichment of proteins involved with regulation of translation, protein folding and mRNA splicing. In approaches based on these data, we found that M2-2 expression downregulates eiF2α phosphorylation and inhibits stress granules assembly under arsenite induction. In addition, we also verified that M2-2 inhibits translation initiation, and is targeted for proteasome degradation, being localized to granules composed by defective ribosomal products at the cytoplasm. These results suggest that besides its functions in the regulation of genome replication, M2-2 may exert additional functions to contribute to successful RSV infection.

**Author summary:** Exploring how viruses take control of the cellular machinery is a common strategy to understand the infection process and to identify targets for inhibition of virus replication. In this work we investigated the cellular functions of the protein M2-2 from the respiratory syncytial virus. Although this virus is an important pathogen responsible for respiratory infections in immunocompromised individuals, currently there are no vaccines or effective treatments to inhibit its infection. Our findings showed that the protein M2-2 interferes with protein synthesis, being able to downregulate the assembly of stress granules during stress stimuli. Besides, we verified the relationship between M2-2 and the proteasome machinery, which is responsible for protein degradation and is also involved with protein synthesis. These results present new functions for the protein M2-2, indicating additional mechanisms utilized by the virus to facilitate infection, providing new perspectives for the search of antiviral targets.

## Introduction

The Respiratory Syncytial Virus (RSV) is an enveloped, single-stranded, negative RNA virus belonging to the order *Mononegavirales* family *Pneumoviridae* [1]. It is one of the major causes of lower tract respiratory disorders, being mainly responsible for infections in newborns, infants, elderly, and immunocompromised individuals. Despite its importance as a pathogen, currently there are no vaccines or effective drugs available against the virus, turning its replicative cycle into an interesting target for the discovery of new antiviral agents [2, 3].

RSV infection starts with the fusion of the viral envelope to the cell membrane and the release of its ribonucleoprotein complex into the cytoplasm. The last comprises the viral nucleocapsid, the RNA-dependent RNA polymerase (RdRp) and its cofactor for transcription. The nucleocapsid is composed by the viral genome associated with the nucleoprotein N, which works as a template for the polymerase while it keeps the genomic RNA hidden [4]. The RdRp has two main components, the phosphoprotein P, a tetrameric protein able to interact with the nucleocapsid and the large protein L, responsible for the catalytic activity of the polymerase [5, 6]. The last performs both transcription and replication activities, which are coordinated by cofactors transcribed from the two open reading frames (ORFs) of the gene M2. The first ORF transcribes for the M2-1 protein, a regulator of transcription and structural component of the virion, while the second ORF transcribes for the M2-2 protein, thought to direct the polymerase activity for replication [7, 8, 9].

Once the ribonucleoprotein complex gets into the cytoplasm, it starts rounds of transcription to produce viral proteins. Few hours post infection, cytoplasmic inclusion bodies (IBs) are assembled, a process triggered by the proteins N and P, able to phase separate when co-expressed [10, 11]. These structures work as viral factories for genome transcription and replication, processes highly regulated by M2-1 and M2-2. M2-1 is essential for viral transcription, and its role on transcription and processing of viral mRNAs has been broadly explored [12, 13]. On the other hand, M2-2 function was firstly deduced from the phenotype produced by a mutant virus lacking its ORF. Infection by this mutant showed enhanced mRNA production with delayed accumulation of viral genome, suggesting a role in switching the polymerase activity for replication [7, 14]. This observation was recently confirmed by Blanchard and colleagues, showing that M2-2 expression can rearrange the interactions between L and other IB components [15].

In addition to its exerted functions in replication, viral proteins are thought to hijack cellular components to assist viral infection or to impair cellular metabolism and host defenses. In this context, RSV proteins have been shown to interfere with many cellular processes. Non-structural proteins NS1 and NS2 prevent IFN I induction, and NS1 is also capable to downregulate the expression of interferon-stimulated genes in the nucleus [16, 17]. In the same way, the matrix protein M also localizes to the nucleus and impairs the expression of genes related to mitochondrial metabolism [18]. On the cytoplasm, N and P hijack cellular chaperones to aid viral replication and impair host defenses by sequestering and keeping innate immune effectors inside of IBs [19, 20]. Accordingly, M2-1 also recruits cellular proteins related to mRNA maintenance to support the synthesis and processing of viral transcripts inside of IBs [13, 21].

Although much has been made to explore the role of RSV proteins in rearranging cellular machinery to its own favor, little has been done to investigate the relationship of M2-2 with cellular components. Previous data on Vero E6 cells showed that M2-2 is poorly localized to IBs, and accumulates in cytoplasmic granules and nucleus, indicating it could have additional functions during infection [15]. To look at this question, in this work we investigated the relation between M2-2 and the cellular machinery. Expressing FLAG-tagged M2-2, we identified by co-immunoprecipitation assay coupled to mass spectrometry analysis a high correlation of M2-2 with ribosomal components, chaperones, splicing factors, and proteasome subunits. Looking for a correlation between proteomic data and M2-2 cytoplasmic granules, we found that M2-2 expression downregulates eiF2α phosphorylation and prevents stress granules assembly. Additionally, we also verified the ability of M2-2 in regulating translation, being able to inhibit both 5’ cap and IRES-dependent translation initiation. Finally, we characterized M2-2 cytoplasmic granules as defective ribosomal products (DRiPs) and showed that the protein is targeted for proteasome degradation, implying it may be involved with translation elongation as well. These data reveal new undescribed functions for M2-2 and suggest it as a multitask protein rather than just a polymerase cofactor.

## Results

### Expression of FLAG and EGFP tagged M2-2 shows similar cellular distribution and partial localization to viral inclusion bodies

As mentioned above, the role of M2-2 on the modulation of the viral polymerase activity has already been shown by expressing a c-myc tagged M2-2 in Vero E6 cells co-infected with RSV [15]. Despite its known role as a polymerase cofactor for genome replication, M2-2 showed only partial recruitment to viral IBs, being mainly localized to nucleus and cytoplasmic granules, suggesting it may be involved in events other than regulating the polymerase activity.

To further explore this possibility, we synthesized two codon-optimized M2-2 genes, tagged by FLAG or EGFP at their amino terminus. Because M2-2 is a small 10 kDa protein, expressing it fused to EGFP could lead to its mislocalization, so we first evaluated the expression of both proteins in HEK293T cells, and their distribution in Hep-2 and Vero E6 cells (Fig 1a and b, S1a Fig). Similar to a previous report [15], both FLAG and EGFP-M2-2 presented nuclear and cytoplasmic localization in a tag-independent way. However, we also noticed cells where M2-2 assumed different patterns of distribution, presenting no cytoplasmic granules, showing nuclear foci or, at a minor amount, getting an ER-like shape, which we later confirmed by co-staining with the ER marker calreticulin (S1b Fig). Despite their non-homogeneity, when co-expressed, FLAG and EGFP-M2-2 were recruited to the same sites, indicating that M2-2 may maintain its interactions regardless of the amino-terminal tags (Fig 1c).

**Fig 1.**
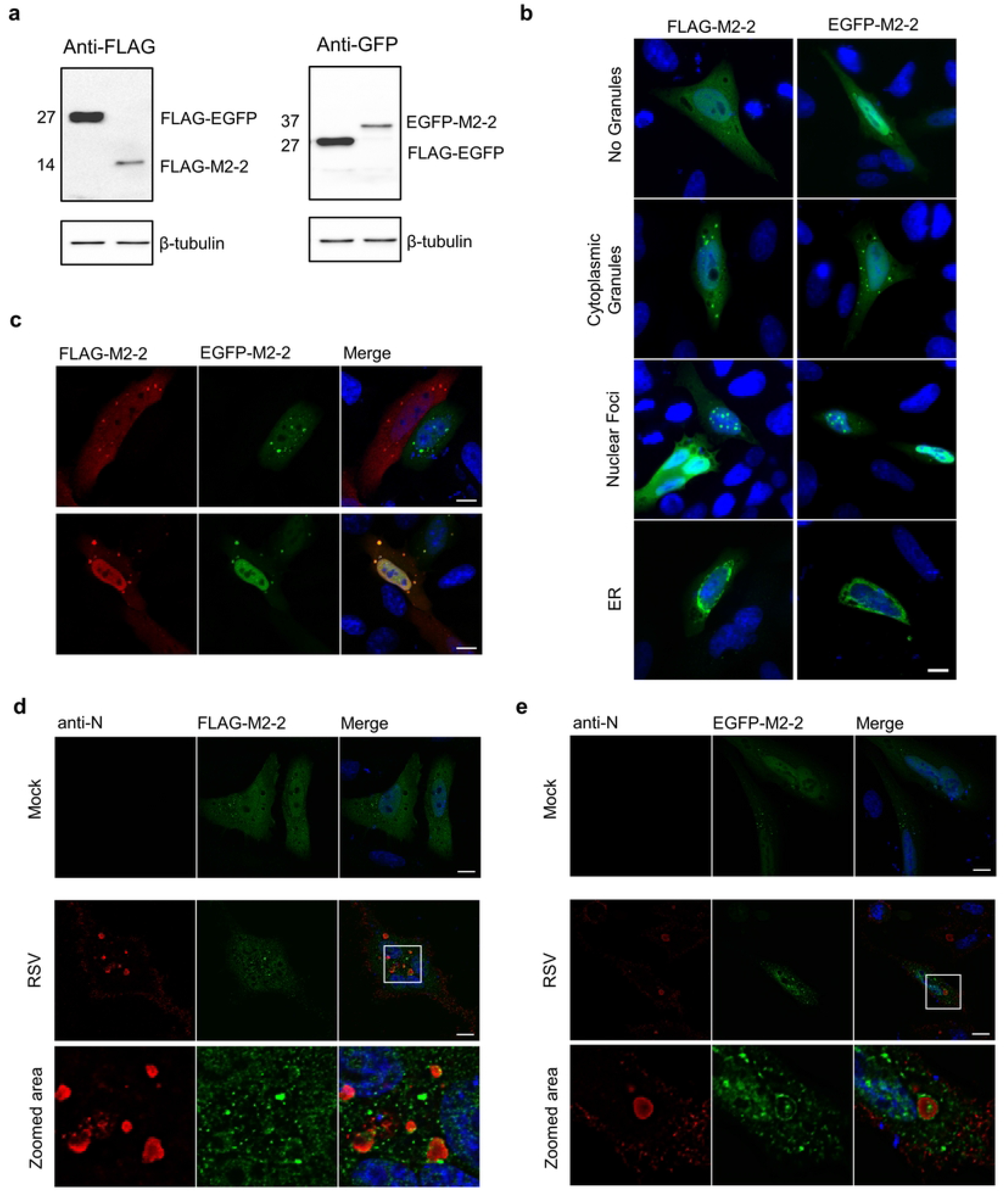
Characterization of recombinant M2-2 expression in human cell lines. (a) Detection of FLAG and EGFP-M2-2 expression in HEK293T cells at 24 hpt. FLAG-EGFP was used as a control for protein detection. Observed molecular weights are shown on the left. (b) Immunofluorescence of FLAG-M2-2 and EGFP-M2-2 expressed for 24h in HEp-2 cells. Panels show distinct distribution patterns observed for both proteins, as indicated on the left. (c) Confocal immunofluorescence showing co-expression of FLAG-M2-2 and EGFP-M2-2 in the same cell (second row) or expressed alone (first row). (d) Expression of FLAG-M2-2 or EGFP-M2-2 (e) in mock or RSV infected HEp-2 cells. Though M2-2 presents nuclear and cytoplasmic diffuse distributions, recruitment to IBs, stained with anti-N, can be observed on zoomed areas. Images were taken with a Zeiss LSM-780-NLO microscope. Raw images (d, e) were submitted to deconvolution as described in the methods section. All images are representative of at least three independent experiments. Scale bars 10 μm.

Finally, we also expressed FLAG and EGFP-M2-2 in infected Hep-2 cells, observing only a partial recruitment of both proteins to viral IBs (Fig 1d and e), reinforcing the previous report obtained in RSV co-infected Vero E6 cells [15]. Together, these data confirm the recruitment of M2-2 to IBs in a human cell line and point to its involvement with the cellular machinery in both nucleus and cytoplasm, either expressed alone or during infection. This makes way to investigate potential M2-2 cellular partners and to explore unidentified functions performed by the protein to enable successful infection.

### Proteomic analysis reveals translation and splicing machineries as potential interactors of M2-2

Aiming to identify potential M2-2 partners, we expressed FLAG-M2-2 or empty vector in HEK293T cells and performed immunoprecipitation (Fig 2a) followed by tryptic digestion and mass spectrometry analysis. Potential interactors identified in at least two of three independent experiments were selected for further analysis, after exclusion of proteins identified on negative control samples. Among the 72 proteins found (Fig 2b, S1 Table), we observed a surprising number of ribosomal proteins, splicing factors and chaperones as possible interactors. Accordingly, we identified enriched Gene Ontology (GO) terms for Biological Process associated to regulation of translation and translation elongation, and terms related to processing, stability and splicing of RNA and mRNA (Fig 2c).

**Fig 2.**
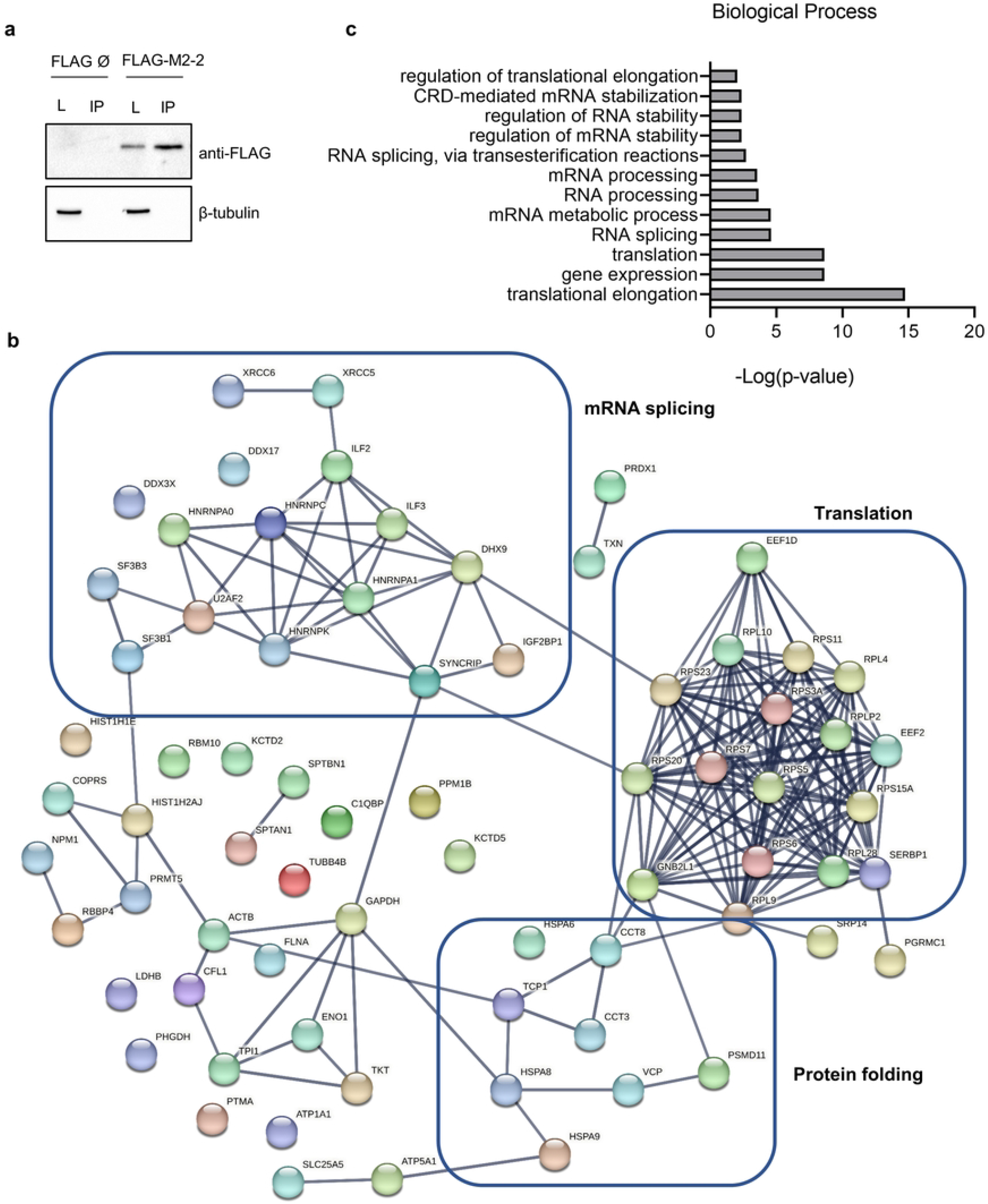
Proteomic analysis of M2-2 interactors shows enrichment of ribosome components, chaperones and splicing factors. (a) FLAG-M2-2 or empty vector were expressed in HEK293T cells for 48h and submitted to immunoprecipitation protocol. Western blot shows detection of FLAG-M2-2 in lysate (L) and eluted proteins (IP). Images are representative of three independent experiments sent for mass spectrometry analysis. (b) STRING network of the 72 proteins identified by mass spectrometry as potential interactors of M2-2. Lines connecting nodes indicate high-confidence interactions, as set during analysis. Protein names are indicated on nodes. Three main clusters are highlighted on the network, showing proteins related to translation, protein folding and mRNA splicing. (c) Graph showing enriched GO terms for Biological Process. Enrichment analysis for the identified proteins was performed on BinGO, as described in the methods section.

These findings are endorsed by our previous observations, since M2-2 is localized to the nucleus, where mRNA synthesis and processing takes place, as well as at the cytoplasm, where it is mostly recruited to cellular granules rather than viral IBs during infection. Stress granules (SGs) and p-bodies are cytoplasmic granules known to be assembled by interactions between proteins and RNA and are described to be target of viral proteins to facilitate replication [22, 23]. Because the proteome of SGs harbor some of the ribosomal proteins, chaperones, and other RNA binding proteins [24] identified as potential partners of M2-2, we decided to investigate its relationship with these structures.

### M2-2 inhibits SGs assembly by keeping a low level of phosphorylated eiF2α

To identify if M2-2 is recruited to SGs, we expressed FLAG and EGFP-M2-2 in Hep-2 cells and performed immunofluorescence using as SG marker the protein G3BP1. As a negative control, cells were treated with the small molecule ISRIB (Integrated Stress Response Inhibitor), known to counteract the inhibitory effect of the phosphorylated eiF2α on the eiF2B complex, promoting disassembly of SGs [25, 26], while for induction of SGs, cells were treated with sodium arsenite, an inducer of oxidative stress. Figures 3a and 3b show that neither G3BP1 is recruited to M2-2 granules, nor ISRIB is able to promote their disassembly. Moreover, arsenite-induced SGs revealed to be distinct structures from the ones carrying M2-2 (Fig 3a and b -zoomed areas). Similar results were obtained with PABP (S2a and b Fig), another SG component [24], or during the co-expression of M2-2 and tagged YB-1 proteins, which are recruited for both SGs and p-bodies [27, 28], showing the absence of M2-2 in these granules as well (S2c and d Fig).

**Fig 3.**
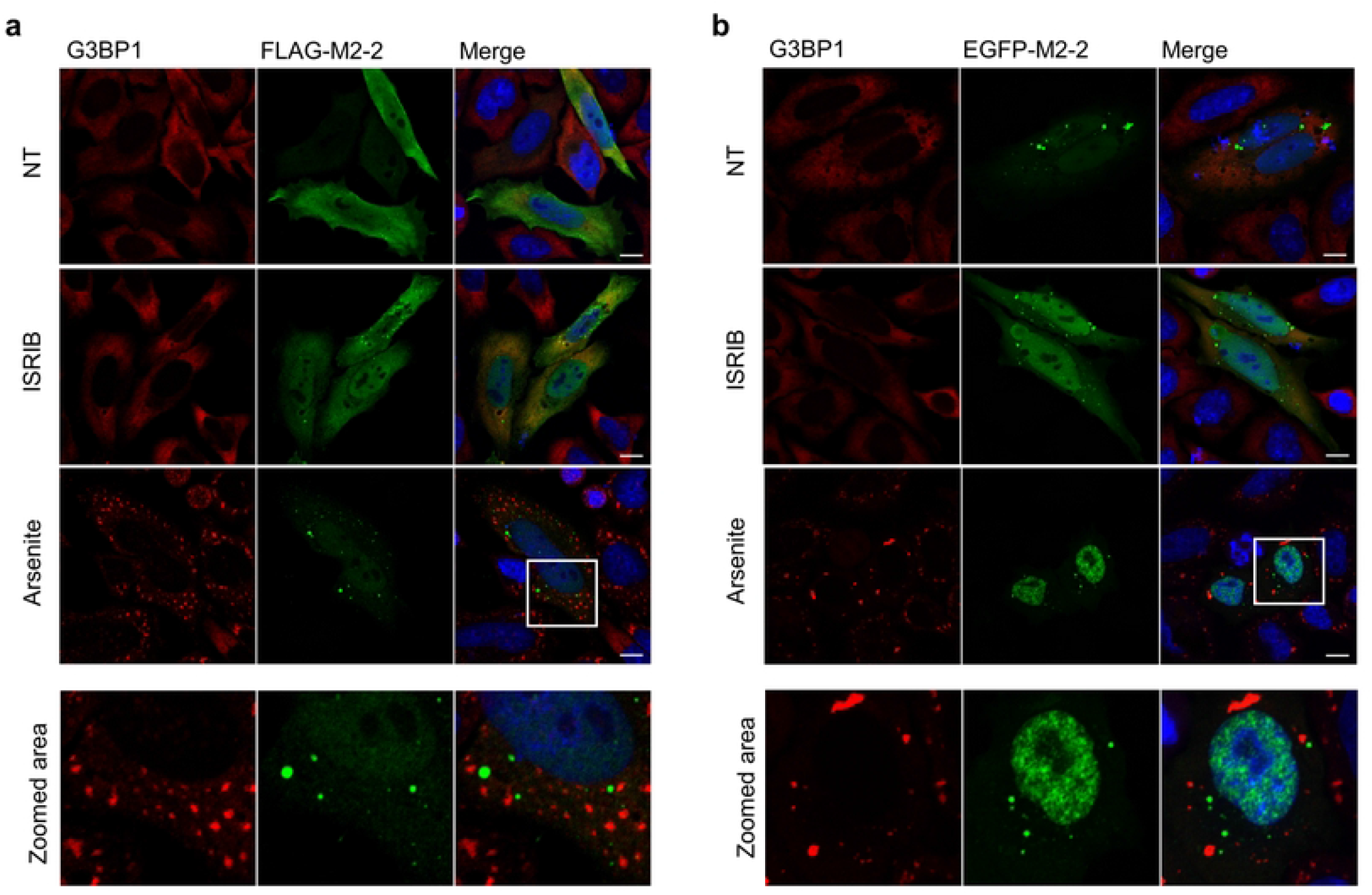
M2-2 does not colocalize with G3BP1 and stress granules. Expression of FLAG-M2-2 (a) and EGFP-M2-2 (b) in HEp-2 cells stained with anti-G3BP1. Cells were treated with 200 nM ISRIB (for 20h) or 0.5 mM arsenite for 30 min before fixation. Non-treated (NT) and ISRIB cells show diffuse G3BP1 distribution, while SGs can be observed under arsenite treatment. Distinction between M2-2 granules and SGs can be seen in zoomed areas. All images were taken with a Zeiss LSM-780-NLO microscope and are representative of three independent experiments. Scale bars 10 µm.

Although no interaction with SGs has been observed, we did notice a portion of cells expressing M2-2 where arsenite was not able to induce their assembly (Fig 4a), with a proportion of roughly 40% over three independent experiments (Fig 4b). Besides, similar inhibition could be seen on A549 and Vero E6 cells (S3a and b Fig). Stress granules are assembled in response to eiF2α phosphorylation [23], therefore we expressed FLAG-M2-2 and EGFP-M2-2 or control vectors in HEK293T cells and evaluated the phosphorylation level of eiF2α, using as negative control transfected cells treated with ISRIB (Fig 4c). Consistent with the above results, cells expressing M2-2 showed a significantly reduced level of phosphorylated eiF2α when compared to empty vector or FLAG-EGFP transfected cells (Fig 4d), implying it may contribute to hamper host defenses coordinated by eiF2α phosphorylation during infection.

**Fig 4.**
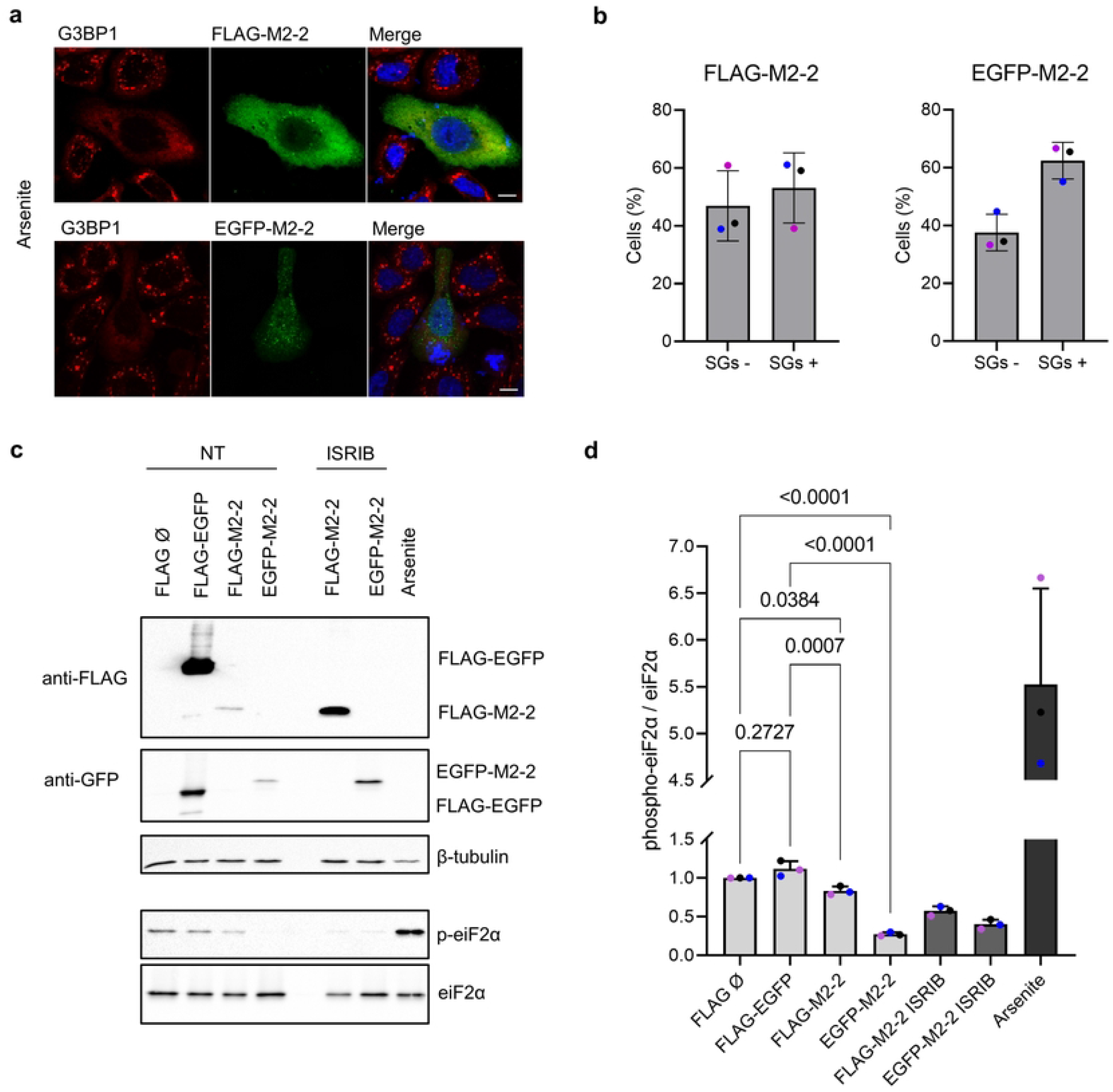
M2-2 expression inhibits SGs assembly and eiF2α phosphorylation. (a) Immunofluorescence of FLAG-M2-2 and EGFP-M2-2 in HEp-2 cells treated with arsenite (0.5 mM for 30 min). Inhibition of SGs assembly, stained by G3BP1, can be seen in cells expressing M2-2. (b) Graphs present the proportion between M2-2 expressing cells positive and negative for stress granules. Roughly 30 cells were counted in each independent experiment, and colored dots indicate individual means for each experiment (n=3). Error bars show standard deviation. (c) FLAG-M2-2 and EGFP-M2-2 or controls (FLAG-EGFP and empty vector) were transfected in HEK293T cells, and after 24h cells were lysed and analyzed by western blot. Treatments with ISRIB (200 nM) or arsenite (0.5 mM) were performed as previously described. Phosphorylation of eiF2α was quantified and submitted to one-way anova followed by Tukey’s multiple comparisons test (d). Colored dots indicate values from independent replicates (n=3). Error bars show standard deviation and p-values are shown in the graph.

Previous reports already explored the relation between RSV infection and the activation of the integrated stress response mediated by eiF2α phosphorylation, showing that infection does not induce stress granules, and that RSV prevents eiF2α phosphorylation by the interaction between N protein and the kinase PKR [29, 30, 30]. In our experiments, RSV infection did not trigger eiF2α phosphorylation in HEK293T, Hep-2 and Vero E6 cells at 24 hpi (S4a Fig). Beyond that, immunofluorescence of Hep-2 cells infected for 24h showed no induction of SGs, and even some inhibition of their assembly under slight arsenite treatment (S4b Fig). These results confirm preceding data for RSV infection and suggest a new role for M2-2 as a stress granule antagonist.

### M2-2 downregulates translation, which is reversed by ISRIB treatment

Regardless of our prior aim in evaluating the phosphorylation of eiF2α, we observed a surprising enhancement in the expression of M2-2 under ISRIB treatment compared to non-treated cells (Fig 4c). Indeed, our initial data showed lower expression levels for both tagged M2-2 proteins when compared to FLAG-EGFP (Fig 1a), however, the mechanism of action described for ISRIB does not include stimulus of basal translation [25, 32]. To ascertain if the effect of ISRIB on M2-2 expression was specific, we expressed FLAG and EGFP-M2-2 in HEK293T cells, using as controls FLAG-M2-1 or FLAG-YB-1 proteins. As expected, ISRIB showed no improvement in the expression of the control proteins over time, contrasting to its effect on M2-2 (S5a and b Fig).

Because ISRIB is related to the initial steps of translation, we decided to examine the effect of M2-2 expression on protein syntheses, which could explain the effect of ISRIB on its own expression levels. Thus, we expressed FLAG-EGFP, or tagged M2-2 proteins in HEK293T cells and 24 hpt (hours post-transfection) we performed a SUnSET assay to assess the translation rate measured by puromycin incorporation [33]. Curiously, both M2-2 transfected cells presented diminished translation when compared to FLAG-EGFP (Fig 5a and b). Additionally, we also co-expressed tagged M2-2 and FLAG-EGFP proteins with a dual luciferase reporter system to evaluate inhibition of different translation initiation mechanisms, containing a bicistronic 5’ cap-dependent Renilla luciferase in tandem with an IRES-dependent Firefly luciferase [34]. Supporting SUnSET results, M2-2 could hamper the expression of both luciferases, with a more prominent effect on cap-dependent translation (Fig 5c).

**Fig 5.**
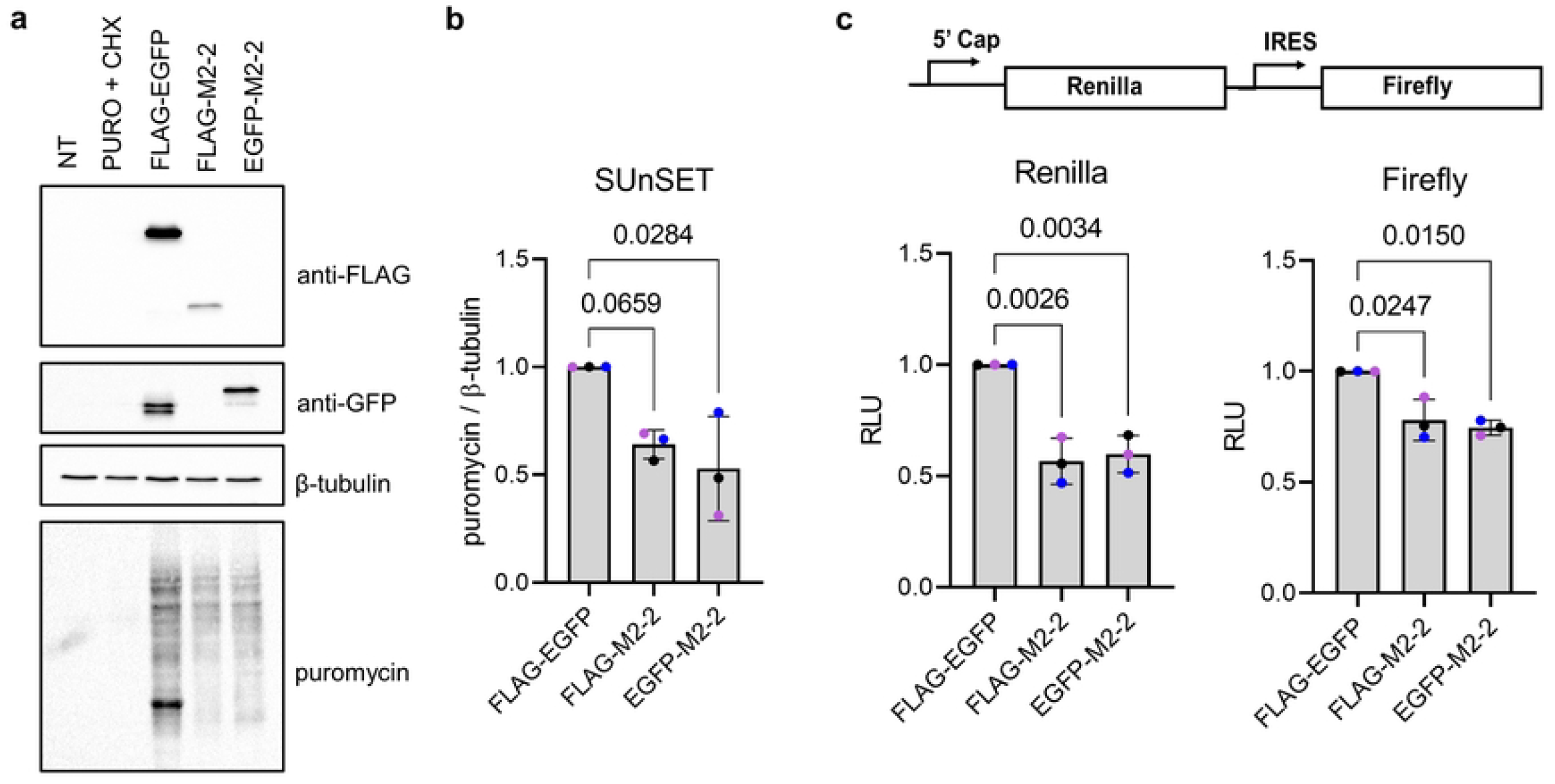
M2-2 expression inhibits translation initiation. (a) SUnSET assay performed in HEK293T cells expressing the control FLAG-EGFP, or FLAG and EGFP-M2-2 proteins. After 24h, cells were incubated with puromycin (10 µg/mL) for 10 min, rinsed and lysed for western blot analysis. Puromycin with cycloheximide (100 µg/mL) or non-treated cells were used as controls. Translation rates, indicated by puromycin staining, are quantified in (b), and all images are representative of three independent experiments (n=3). (c) Reporter luciferase assay evaluating the inhibitory effect of M2-2 in the translation of Renilla (5’ cap) and Firefly (IRES) luciferases, as shown above. Graphs show relative luminescence units (RLU) obtained in three independent experiments performed in triplicate (n=3). Colored dots shown in (b) and (c) indicate individual means obtained in each individual experiment. Statistical analysis was performed by one-way anova followed by Dunnett’s multiple comparisons test. Error bars indicate standard deviation and p-values are indicated on the graph.

Except for the identification of RACK1 (also identified as GNB2L1), described as a regulator of translation initiation [35, 36], the other proteins identified by proteomic analysis (Fig 2b, S1 Table), though related to translation, are not described as main components of translation initiation, nor part of the path regulated by eiF2B – eiF2 ternary complex – 43S preinitiation complex, which is a target of ISRIB [37]. Besides, analysis of enriched GO terms pointed translation and translation elongation, but not translation initiation, as functions performed by M2-2 potential partners (Fig 2c). Even if these findings are not totally in agreement, our presented results reveal a new role for the M2-2 protein as an inhibitor of translation.

### RSV infection inhibits 5’ cap and IRES-dependent translation without affecting global protein synthesis

To find out if RSV infection could drive inhibitory effects on protein translation similar to M2-2 expression, we infected HEK293T cells with RSV, and 1 hpi (hour post-infection) cells were transfected with the plasmid for expression of the reporter luciferases. Graphs in Supplementary Figure 6a show reduced luciferase activity for both promoters in infected cells, resembling that of M2-2. Hoping to see similar results, we also infected HEK293T, Hep-2 and Vero E6 cells, and performed SUnSET assay at 24h, 36h and 48hpi. Unexpectedly, puromycin incorporation was not suppressed during infection, indicating even higher translation rates in Hep-2 and Vero E6 cells (S6b and c Fig).

**Fig 6.**
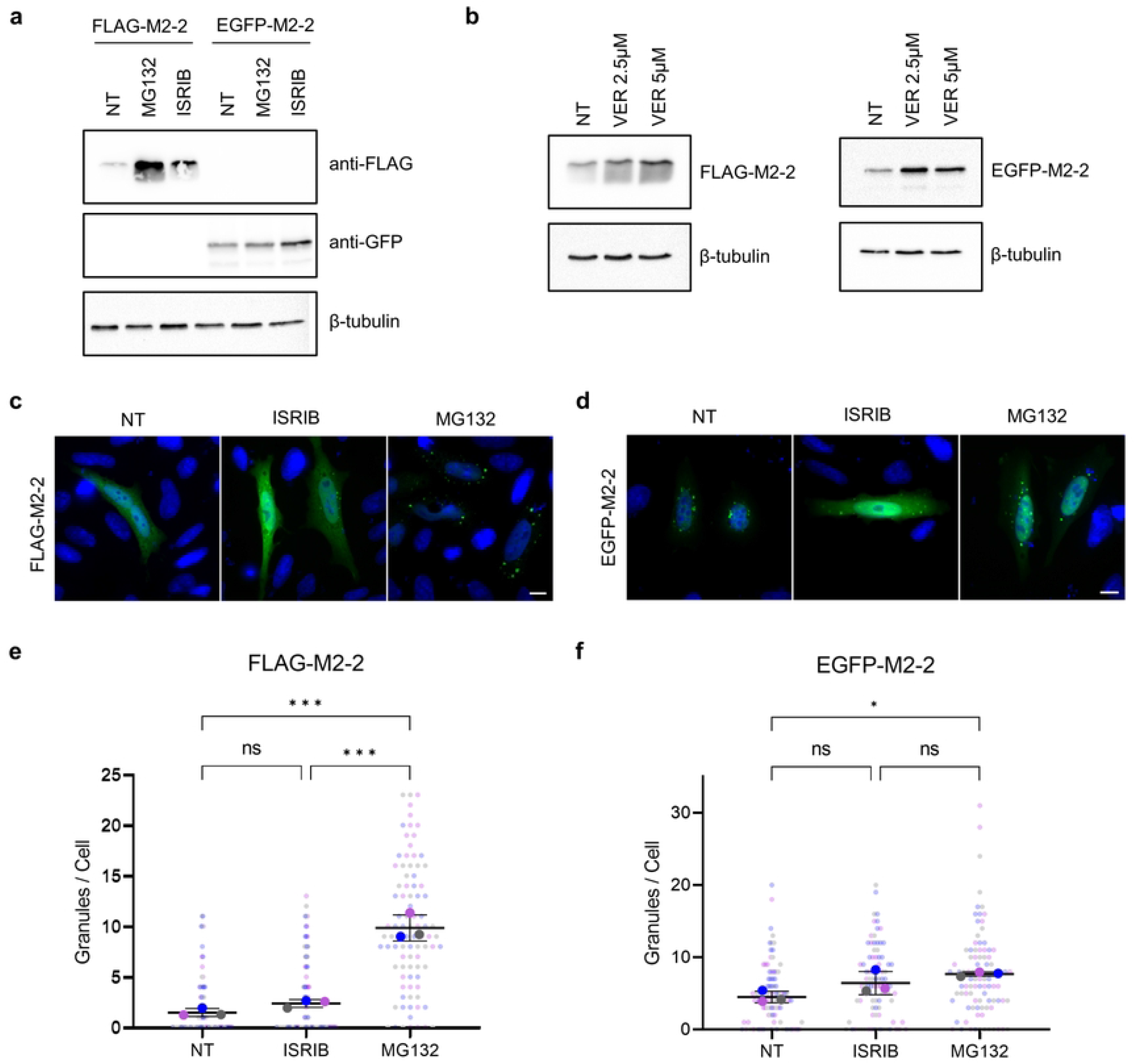
Proteasome inhibition promotes FLAG-M2-2 phase separation and increases its detection by western blot. (a) Western blot detection of FLAG-M2-2 and EGFP-M2-2 expressed for 24h in HEK293T cells non-treated (NT), or under ISRIB (200 nM) and MG132 (5 µM) treatments. (b) Expression of FLAG and EGFP-M2-2 under VER (NT, 2.5 or 5 µM) treatment as in (a). Western blot images in (a) and (b) are representative of three independent experiments. Cellular distribution of FLAG-M2-2 (c) and EGFP-M2-2 (d) under different treatments (ISRIB 200 nM for 20h, MG132 5 µM for 2h) in HEp-2 cells, showing increase in the number of granules stained by FLAG-M2-2 under MG132 treatment. (e) and (f) present the average number of granules by cell for each protein submitted to different treatments. Small dots show individual number of counted granules over 30 cells for each condition, while bigger dots present mean values. Different colors show values obtained across each independent experiment (n=3). Statistical differences were evaluated by one-way anova following Tukey’s multiple comparisons test. Error bars indicate standard deviation, and p-values are indicated by * p < 0.05 and *** p < 0.001, ns = not significant.

In the course of infection, many viruses take control of the cellular machinery to facilitate the expression of their own proteins. Since RSV transcripts are flanked by different 5’ UTR sequences, one possibility is that the virus can favor its protein production while it hampers translation of cellular transcripts, a known mechanism described for some cellular proteins and different viruses [38, 39, 40]. Accordingly, though infection has been shown to inhibit 5’ cap-dependent and IRES-dependent translation of reporter luciferases, SUnSET assay may not be suitable to revalidate its inhibition, seeing that viral proteins are still being highly expressed.

### M2-2 is targeted for proteasome degradation and is directed to cytoplasmic granules during MG132 treatment

In the above results, we described two unpredicted roles for M2-2, modulating translation and the cellular response to stress through eiF2α phosphorylation. Nevertheless, these functions do not directly correlate to proteomic data obtained, nor were we able to identify the granular structures to which M2-2 is recruited. Because many of the proteins identified by co-IP were mainly associated with translation elongation, we chose to investigate potential paths connecting it with viral infection, raising a possible association of M2-2 with ribosome quality control (RQC), among other mechanisms involved with this step of translation [41, 42, 43]. M2-2 potential partners encompass, beyond ribosomal proteins, numerous chaperones, including VCP, recently described as necessary for proper RSV infection [44]. Moreover, besides the already described RACK1, we also detected the proteasome subunit PSMD11, all known proteins whose role in RQC has been for long described [45, 46, 47, 48].

In the course of peptide elongation, stalled ribosomes or incorrect folding of the newly synthesized proteins can signal for RQC effectors to ubiquitinate and dismount 80S ribosomal subunit, addressing nascent peptides for proteasome degradation, a process assisted by accessory chaperones [45, 46, 49]. In an attempt to find out whether M2-2 was involved with this process, HEK293T cells were transfected with FLAG and EGFP-M2-2 and treated with the proteasome inhibitor MG132, using as controls ISRIB or non-treated cells. MG132 showed a modest effect on EGFP-M2-2 expression, whereas a surprising improvement could be seen in FLAG-M2-2 detection (Figs 6a and 7d, S9a and 10a Figs), with similar effects observed in Hep-2 cells (S7a Fig). Alternatively, HEK293T cells were treated with the HSP70 inhibitor VER, achieving comparable results (Fig 6b).

**Fig 7.**
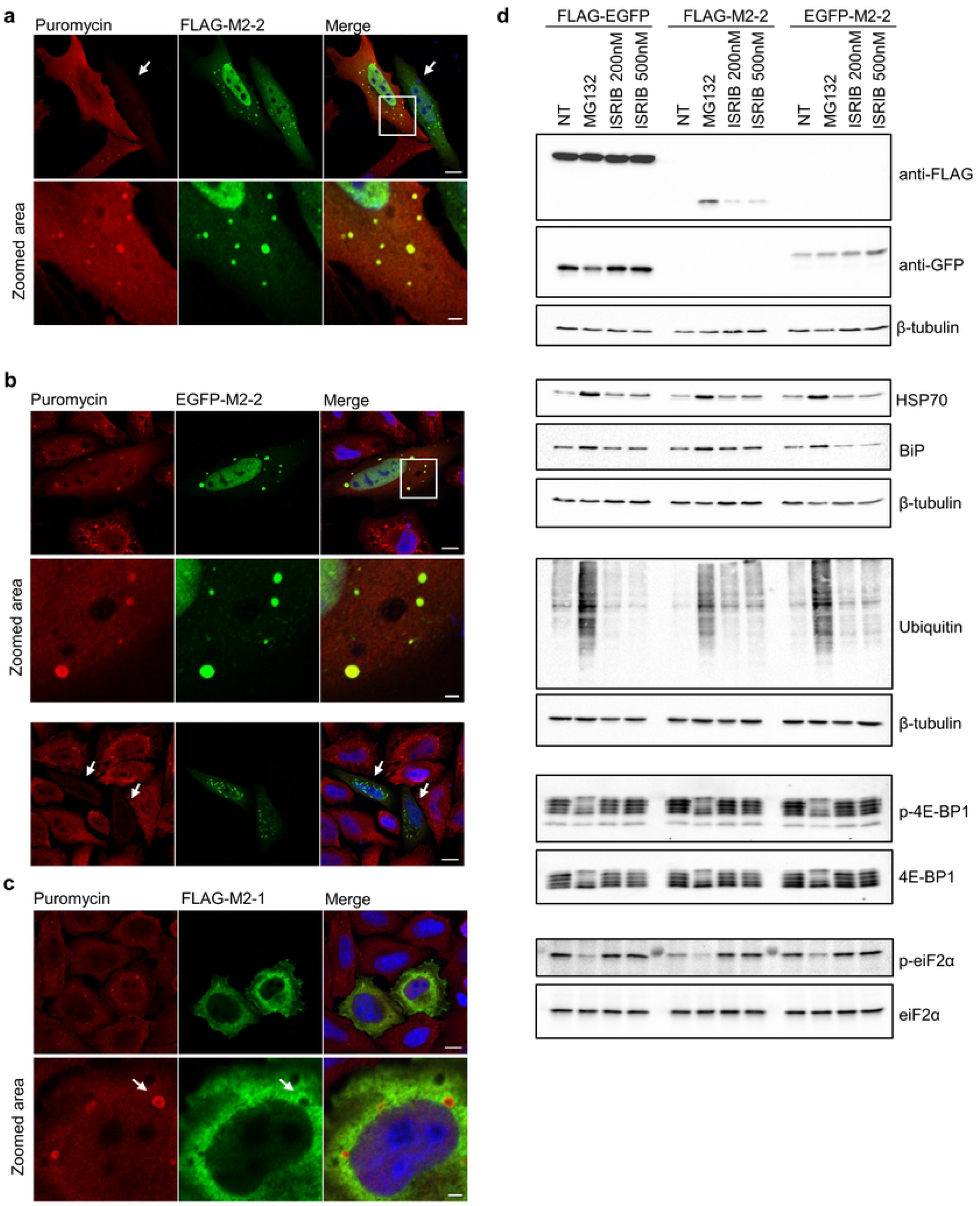
M2-2 interacts with defective proteins targeted for degradation but does not trigger proteostatic stress. FLAG-M2-2 (a) and EGFP-M2-2 (b) were expressed in HEp-2 cells for 24h, and then cells were incubated with puromycin (10 µg/mL) and MG132 (5 µM) for 2h. After treatment, cells were rinsed, fixed, and stained with anti-puromycin for tracking of DRiPs, showing colocalization with M2-2. Inhibition of translation can be observed in cells presenting low puromycin incorporation (white arrows in (a) and (b) – third row). (c) As a control, FLAG-M2-1 was expressed and submitted to the same treatments. White arrows in the zoomed area show absence of colocalization between puromycin stained granules and FLAG-M2-1. All images were taken with a Zeiss LSM-780-NLO microscope and are representative of three independent experiments. Scales bars are 10 µm for full size panels, and 2 µm for zoomed panels. (d) FLAG-EGFP or FLAG and EGFP-M2-2 proteins were expressed in HEK293T under the following treatments, NT – non-treated, MG132 (5 µM), ISRIB (200 nM or 500 nM). After 24h, cells were lysed and extracts were analyzed by western blot, looking for markers of proteostatic stress (HSP70 and BiP), as well as regulators of translation initiation (phospho-4E-BP1 and phospho-eiF2α). Ubiquitin levels were also accessed for evaluation of proteasome inhibition by MG132. Images are representative of three independent experiments.

In addition to western blot data, we verified that MG132, but not ISRIB, is able to induce phase separation of FLAG-M2-2 into cytoplasmic granules in either Hep-2 or A549 cells, whereas only a minor effect is observed for EGFP-M2-2 (Fig 6c - f, S7b Fig). However, when trying to reproduce these data in Vero E6 cells, we could not observe either the improvement in FLAG-M2-2 expression or the increase in the number of its cytoplasmic granules (S7c - e Fig). These results suggest that, in human cells, M2-2 is targeted for proteasomal degradation, which may be prevented by the fusion of EGFP on its amino terminus. It also explains why EGFP-M2-2 displays a slightly more granular pattern than FLAG-M2-2 (Fig 6e and f), given that FLAG-M2-2 granules must be more efficiently cleared by proteasome activity.

### M2-2 colocalizes with granules composed of defective ribosomal products

It has already been described that inhibition of proteasome could trigger aggregation of ubiquitinated peptides addressed for proteasome-dependent degradation, which can be tracked by puromycin-induced termination of translation [50, 51]. Therefore, Hep-2 cells were transfected with FLAG-M2-2 or EGFP-M2-2, and 24 hours post-transfection cells were treated with puromycin and MG132 for 2h. As a negative control, cells were transfected with FLAG-M2-1. Figure 7 shows colocalization between puromycin granules and both M2-2 tagged proteins, contrasting to FLAG-M2-1, for which colocalization was absent (Fig 7a – c). Furthermore, we could detect cells where puromycin incorporation was inhibited (Fig 7a and b – white arrows), confirming our preceding results for inhibition of translation. These results indicate that M2-2 is involved in different steps of translation, inhibiting its initiation, and interacting with newly synthesized peptides targeted for degradation. However, if M2-2 itself promotes elongation arrest or premature termination of translation, remains to be cleared.

Despite the discrepant results regarding the effect of proteasome inhibition in Vero E6 cells, FLAG and EGFP-M2-2 also exhibited colocalization with puromycin granules in this cell line, as well as inhibition of protein synthesis evidenced by low puromycin incorporation (S8a and b Fig). Concerning these results, we point out that even if both proteins are still able to inhibit translation and colocalize with puromycin-stained granules in this lineage, Vero E6 cells are from a different species, which can result in discrete changes in the cellular proteome [52] and, consequently, in the group of proteins to which M2-2 interacts, leading to variations in the observed phenotype, as the one we have seen for MG132 treatment.

### M2-2 modulates translation without triggering proteostatic stress

Finally, we evaluated whether M2-2 involvement with translation and proteasome machineries could modulate the expression of other related proteins. To do so, we expressed FLAG-EGFP or M2-2 proteins in HEK293T cells under no treatment, or in the presence of ISRIB and MG132. Quantitative analysis showed no differences in the expression level of HSP70 and BiP (Fig 7d, S9b Fig), indicating that the inhibitory effect of M2-2 does not trigger the unfolded protein response [50]. Comparison of ubiquitin levels between ISRIB and non-treated cells also suggests it does not affect proteasome activity, reinforcing that MG132 and ISRIB regulate M2-2 expression by different mechanisms (Fig 7d, S10b Fig). Lastly, in addition to reduced eiF2α phosphorylation, we noticed an enhanced level of the phosphorylated 4E-BP1 protein (Fig 7d, S7b Fig), which contrasts with the observed inhibition of translation initiation [37].

These results show that, though M2-2 accumulates in granules composed of defective proteins, it does not induce a folding stress response. Additionally, the observed levels of phosphorylated eiF2α and 4E-BP1 do not correlate with the inhibition of translation initiation promoted by M2-2, suggesting that these proteins are modulated by a cellular compensatory effect, rather than a direct involvement with M2-2.

## Discussion

In this work, we investigated the yet unexplored relationship between the RSV M2-2 protein and the cellular machinery. In our initial experiments, the M2-2 interactome showed a substantial correlation with the regulation of translation and protein folding. This was later confirmed by the ability of M2-2 to inhibit translation initiation, as well as the integrated stress response, mediated by the phosphorylation of eiF2α and SGs assembly. Furthermore, we also verified that M2-2 is targeted for proteasome degradation, and colocalizes with defective ribosomal products, suggesting it could be involved with translation elongation as well. To our knowledge, this is the first study to describe additional functions performed by M2-2 to modulate the cellular metabolism, indicating a new role for this protein during RSV infection.

In our proteomic analysis, we identified 72 potential cellular partners of M2-2, composed by three main functional clusters, related to mRNA splicing, translation and protein folding (Fig 2b and c, S1 Table), in accordance with our observations exploring the cellular distribution of M2-2 (Fig 1b, S1a Fig). In the nucleus, we detected M2-2 diffusely in the nucleoplasm or in a punctate pattern, resembling what is observed for some splicing effectors [53, 54]. Other RSV proteins have been described to localize into the nucleus acting as inhibitors of transcription [17, 18], with no data exploring their direct relation with mRNA splicing. However, if M2-2 is recruited to sites where splicing takes place, and if it interferes with pre-mRNA processing, remains to be addressed.

In the cytoplasm, M2-2 presented both diffuse and granular distributions, being only partially recruited to viral inclusion bodies (Fig 1d and e). Such a pattern was similar to previous reports exploring the relationship between viral IBs and stress granules [29, 30, 30]. Because some of the M2-2 partners are also described as SG components [24], and considering our observations of its granules as being distinct from IBs, we sought to investigate the relationship between both structures by immunofluorescence. However, we did not detect colocalization between SG markers and M2-2 granules, nor the treatment with ISRIB was able to promote their disassembly (Fig 3a and b). Besides, arsenite-induced SGs presented a different distribution from the granules harboring M2-2. Despite their distinguished distributions, we noticed a significant proportion of cells where M2-2 expression impaired SGs assembly (Fig 4a and b, S3a and b Fig). To ascertain if this observation was related with the main effector for induction of SGs and block of translation, we analyzed the phosphorylation level of eiF2α in cells expressing M2-2. In these cells, we have found phospho-eiF2α substantially reduced in comparison to controls (Fig 4c and d), corroborating the immunofluorescence results. Likewise, RSV infection did not trigger eiF2α phosphorylation in three different cell lines analyzed at 24 hpi (S4a Fig). Additionally, we could not detect phase separation of G3BP1 in HEp-2 infected cells, with even some inhibition of SGs assembly under arsenite treatment (S4b Fig). Though RSV antagonism to SGs has been previously explored [29, 30], it does not exclude M2-2 as one of its effectors.

While the above results suggested a new role for M2-2 as an antagonist of the integrated stress response, we were surprised by the improved expression of M2-2 under ISRIB treatment. ISRIB acts on the eiF2B – eiF2 ternary complex – 43S preinitiation complex branch [25, 32, 37], preventing the inhibitory effect of the phosphorylated eiF2 α-subunit on the eiF2B guanine exchange factor, restoring basal levels of translation, but not stimulating overexpression of proteins [25]. Accordingly, we verified the specificity of ISRIB in M2-2 expression in comparison to control proteins, indicating that M2-2 could be interfering with its own translation (S5a and b Fig). This was verified by SUnSET assay, showing reduced translation rates measured by puromycin incorporation, as well as co-expression of M2-2 with a luciferase reporter system, showing that it was able to inhibit both 5’-cap and IRES-dependent translation initiations (Fig 5a – c).

Endorsing these results, we observed similar inhibition of reporter luciferases during infection, however, we were not able to detect differences in puromycin incorporation between mock and RSV infected cells (S6a – c Fig). Indeed, even under events where translation is inhibited, specific cellular mRNAs are still transcribed [39], which is also observed for many viruses, for which protein production is expected to be elevated [38, 40]. In these viruses, mRNAs containing specific 5’ UTR sequences allow the continuous expression of viral proteins, whereas the translation of cellular mRNAs is hindered. Since transient M2-2 expression under CMV promoter carries ordinary 5’ UTR, it could explain its downregulated expression. However, it remains to be elucidated whether the expression of M2-2 flanked by RSV 5’ UTR presents normal expression levels in the absence of ISRIB.

Finally, looking for a direct correlation between proteomic data and M2-2 distribution into granules, we verified that M2-2 is targeted for proteasome degradation in human cells, which can be inhibited by treatments with MG132 or the HSP70 inhibitor VER (Fig 6a and b, S7a Fig). Besides, we could detect improved phase separation of FLAG-M2-2 when proteasome activity was inhibited, suggesting that M2-2 granules were cleared by this degradation path (Fig 6c – f, S7b Fig). Compared to FLAG-M2-2, EGFP-tagged M2-2 exhibits a more granular pattern at the cytoplasm, indicating its granules are not properly cleared. In agreement, EGFP-M2-2 expression showed only a modest effect under MG132 treatment. Seeing that M2-2 is a small 10 kDa protein, fusing it to EGFP could interfere with some of its interactions, therefore preventing its association with proteasome effectors.

Literature reports describe the accumulation of cytoplasmic granules composed by defective products of translation when proteasome activity is inhibited [50, 51]. To examine if M2-2 was recruited to these structures, we induced premature termination of translation by puromycin treatment, and tracked labeled peptides by immunofluorescence. Surprisingly, puromycin granules showed colocalization with both FLAG and EGFP-M2-2 (Fig 7a and b, S8a Fig). As M2-2 was shown to depend on both proteasome and HSP70 activities to be degraded, we asked if its involvement with DRiPs could stimulate the unfolded protein response. However, we did not see significant changes in the expression of UPR markers, like HSP70 or BiP (Fig 7d, S9b Fig).

Together, these results propose a new function for M2-2 as a translation regulator (Fig 8). During the initial steps of translation, M2-2 showed inhibitory activity, blocking 5’ cap and IRES-dependent protein syntheses, which was reproduced in infected cells. This inhibition was independent of eiF2α phosphorylation or SGs assembly. In addition, we also detected increased levels of phosphorylated 4E-BP1 (Fig 7d, S9b Fig), a marker of active translation [37], even during the inhibition of protein synthesis promoted by M2-2. It suggests that, though M2-2 can counteract SGs induction, eiF2α and 4E-BP1 phosphorylation are modulated in response to low translation rates (Fig 8a), rather than by direct intervention of M2-2 in these proteins or their kinases.

**Fig 8.**
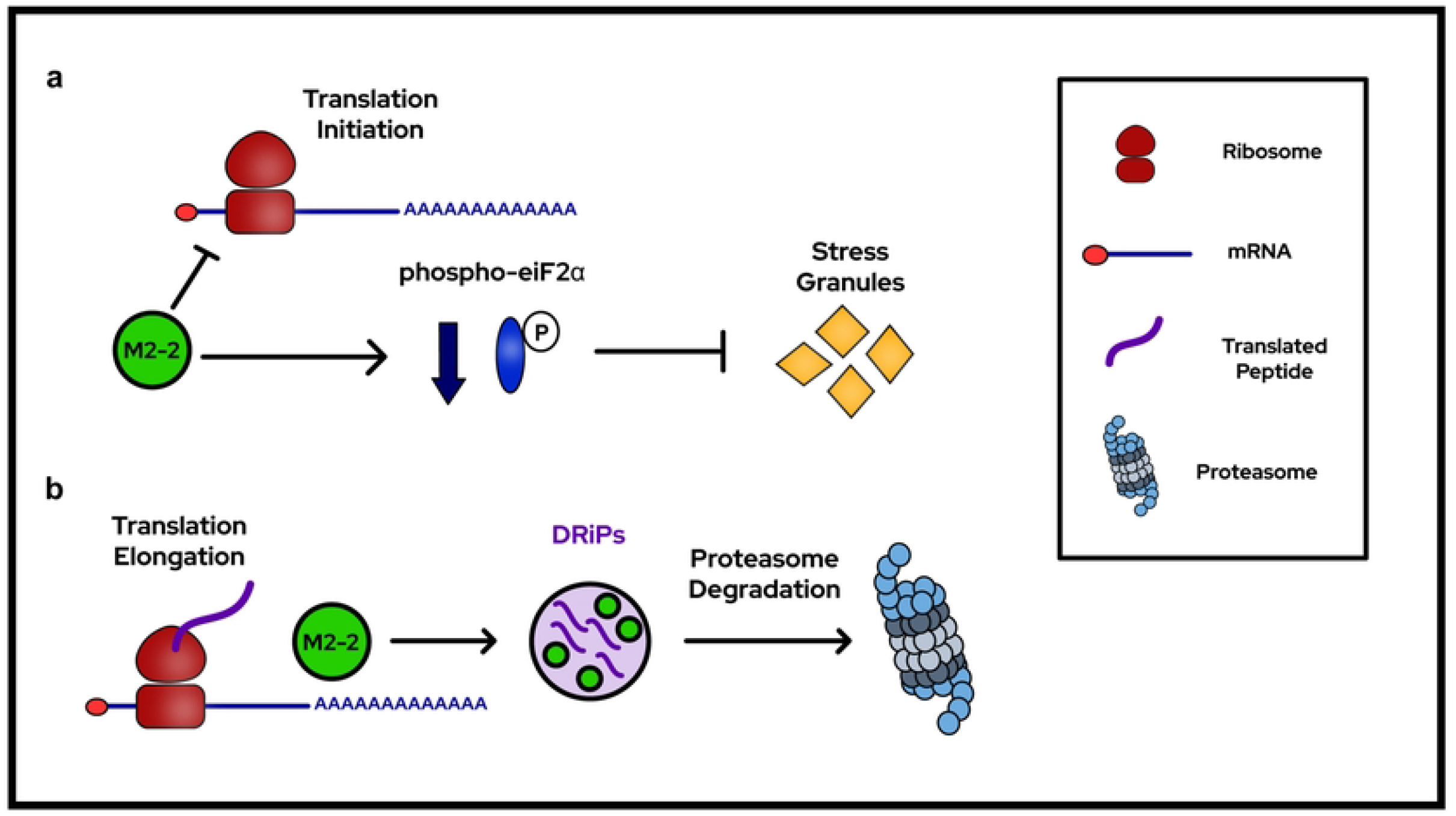
A model describing the involvement of M2-2 with different steps of translation. (a) M2-2 expression inhibits the initial steps of translation. By keeping a reduced translation rate, M2-2 downregulates the phosphorylation of eiF2α, preventing the assembly of stress granules. (b) During the elongation of newly synthesized peptides, M2-2 interacts with defective proteins targeted for degradation. These proteins accumulate into cytoplasmic granules (DRiPs) that are later cleared by proteasome degradation. Figure legend is shown on the right.

On the other hand, M2-2 interaction with DRiPs, as well as its association with the proteasome machinery, implies it could be involved with translation elongation, directing prematurely terminated peptides for degradation (Fig 8b). Such events occur during ribosome quality control activation, where stalled ribosomes are ubiquitinated and dismounted in response to errors in translation or peptides not reaching proper folding, a path utilized by some viruses to favor translation of its own transcripts [41, 42, 43]. Moreover, RQC was also described as an eiF2α antagonist [55], and the premature termination of elongation could contribute for the observed inhibition results obtained in both SUnSET and reporter luciferase assays. In line with this hypothesis, though RSV infection prevents SGs assembly, knockdown of G3BP1 was able to impair viral replication [30], which could be explained by the reduction in the recycling of ubiquitinated ribosomes after RQC induction in these cells [56]. Regardless of these possibilities, whether M2-2 expression or RSV infection can actively promote RQC, remains to be addressed.

Although minor variations were observed in Vero E6 cells (S7c – e Fig), the presented results were consistent between different human cell lines, indicating that care must be taken when characterizing proteins in models from different species. We also point out as a limitation the transient expression of the protein, raising the necessity of further experiments to validate our results with M2-2 expressed during infection. Despite that, this is the first study to explore additional functions for the RSV M2-2 protein, suggesting new mechanisms employed by the virus to subvert the cell machinery to its own favor.

## Materials and Methods

### Cells and virus

HEK293T, HEp-2, A549 and Vero E6 cells were grown at 37°C in Dulbecco’s Modified Eagle Medium (DMEM) supplemented with 10% (v/v) fetal bovine serum (FBS), gentamicin (10 µg/mL) and tylosin (8 µg/mL). Cells were routinely tested for mycoplasma contamination. RSV strain A2 was amplified in HEp-2 cells at 37°C and after two passages, cells were scraped, resuspended in PBS containing 10 mM EDTA (pH 8.0) and Dounce homogenized. Lysates were then centrifuged for precipitation of cell debris (3,250 x g for 20 min at 4°C), and NT buffer (150 mM NaCl, 50 mM Tris–HCl, pH 7.5) containing 50% polyethylene glycol 6000 (w/v) was added to supernatants to a final concentration of 10%. Lysates were incubated for 90 min under gentle agitation (4°C) and then centrifuged (3,250 x g for 20 min at 4°C). Pellets containing purified virus were resuspended in NT buffer containing 15% sucrose (w/v) 1 mM EDTA and stored at -80°C. Plaque assays for stock titration were performed at 37°C in HEp-2 cells, using carboxymethylcellulose (Sigma) as overlay.

### Plasmids

Optimized genes for expression of FLAG-M2-2, EGFP-M2-2, FLAG-YB-1 and EGFP-YB-1 (S1 File) in human cells were synthesized and subcloned in the vector pcDNA3.1(+) at GeneArt (Thermo Fisher Scientific). FLAG-M2-2 gene was inserted between the NcoI and AflII restriction sites, while the other three genes were inserted between NheI and ApaI sites. The optimized gene for M2-1 (S1 File) expression was acquired at GeneArt (Thermo Fisher Scientific), subcloned downstream to FLAG in the vector pcDNA3.1-FLAG [57] (between the BamHI and EcoRI sites) and analyzed by sequencing. Optimization was performed utilizing amino acid sequences obtained on Uniprot (RSV strain A2 for viral genes and *Homo sapiens* for YB-1 protein), and the nucleotide sequence of the optimized genes is provided in the Supplementary Information. Additionally, pcDNA3.1-FLAG (empty vector) and pcDNA3.1-FLAG-EGFP were utilized as control vectors. The vector pLPL, used for the luciferase reporter assay, was previously described [34].

### Transfection and infection

Transfection was performed with Lipofectamine 3000 (Thermo Fisher Scientific) following manufacture’s recommendations. Briefly, cells were transfected with 0.3 µg of DNA for each cm^2^ (transfection surface area), using lipofectamine at the proportion 1:3 (DNA / Lipofectamine) in Opti-MEM (Thermo Fisher Scientific), and after 4 hours, medium was changed for DMEM 1% FBS. For infection, cells were incubated with RSV for 1h at 37°C in DMEM containing 1% FBS, and after adsorption, medium was changed. Transfection for expression of M2-2 plasmids in infected cells was performed 10 hours post-infection, as previously described [15]. For the luciferase reporter assay, cells were transfected 1 hpi. For all experiments, cells were plated the day before to an expected confluence of 70 to 80%.

### Reagents and treatments

ISRIB (SML0843 – Sigma) diluted in DMSO was used at concentrations of 200 nM and 500 nM, and treatments were performed after a change of transfection media (4 hpt until cell lysis or fixation). Arsenite (S7400 – Sigma) diluted in sterile water was used at concentrations of 0.5 mM for 30 min in HEK293T, HEp-2 and A549 cells, and 1 mM for 1h in Vero E6 cells. Puromycin was used at the final concentration of 10 µg/mL, and was incubated for 10 min for SUnSET assays, using co-treatment with cycloheximide (100 µg/mL) as a negative control [33]. For immunofluorescence, cells were incubated with puromycin (10 µg/mL) and MG132 (Abcam) (5 µM) for 2h. Additionally, MG132 treatments were performed for 2h (immunofluorescence) and overnight (4 hpt until cell lysis – for western blot) at 5 µM. The HSP70 inhibitor VER-155008 (Sigma) was used at concentrations of 2.5 µM and 5 µM, with similar treatments to ISRIB.

### Antibodies

The following antibodies were used. Anti-FLAG mouse (F1804 – Sigma, 1:200 IF, 1:2000 WB), anti-FLAG rabbit (F7425 – Sigma, 1:200 IF, 1:2000 WB), anti-GFP rabbit (sc-8334 – Sta Cruz, 1:100 IF, 1:1000 WB), anti-N mouse (ab94806 – Abcam, 1:200 IF), anti-P mouse (ab94965 – Abcam, 1:2000 WB), anti-G3BP1 rabbit (#61559 – Cell Signaling, 1:200 IF), anti-Calreticulin rabbit (ab2907 – Abcam, 1:100 IF), anti-Phospho-eiF2α rabbit (#3398 – Cell Signaling, 1:1000 WB), anti-eiF2α rabbit (#5324 – Cell Signaling, 1:2000 WB), anti-β-tubulin rabbit (ab6046 – Abcam, 1:5000 WB), anti-PABP mouse (sc-32318 – Sta Cruz, 1:10 IF), anti-HSP70 mouse (H5147 – Sigma, 1:2000 WB), anti-BiP rabbit (#3183 – Cell Signaling, 1:1000 WB), anti-Ubiquitin rabbit (#3369 – Cell Signaling, 1:1000 WB), anti-Puromycin mouse (MABE343 – Sigma, 1:200 IF, 1:5000 WB), anti-GAPDH rabbit (ab9485 – Abcam, 1:2000 WB), anti-Phospho-4E-BP1 rabbit (#2855 – Cell Signaling, 1:1000 WB), anti-4E-BP1 rabbit (#9452 – Cell Signaling, 1:1000 WB), anti-Mouse HRP (ab97023 – Abcam, 1:5000 WB), anti-Rabbit HRP (ab97051 – Abcam, 1:5000 WB), anti-Mouse Alexa Fluor 594 (A11032 – Invitrogen, 1:1000 IF), anti-Rabbit Alexa Fluor 488 (A11034 – Invitrogen, 1:1000 IF), anti-Rabbit Alexa Fluor 594 (A11012 – Invitrogen, 1:500 IF).

### Co-immunoprecipitation

HEK293T cells at 70% confluence were transfected with FLAG-M2-2 or empty vector and after 48h cells were rinsed and lysed for 15 min on ice (50 mM Tris-HCl pH 7.5, 150 mM NaCl, 1 mM EDTA, 1% Triton X-100 (v/v), protease and phosphatase inhibitors (Sigma). Lysates were then centrifuged (5,000 x g for 10 min at 4°C) and supernatants incubated with FLAG-beads (Sigma) under gentle agitation (overnight at 4°C), saving a small sample for posterior analysis. Next steps were performed following manufacture’s recommendations. Co-immunoprecipitated proteins were eluted for 1h at 4°C with 3X FLAG peptide (Sigma) in TBS (pH 7.4). Samples were then quantified by BCA assay and analyzed by western blot.

### Sample Preparation and mass spectrometry analysis

Eluted proteins from co-IPs were submitted to trypsin digestion. Briefly, proteins were reduced with Dithiothreitol (DTT) to a final concentration of 10 mM, incubated for 45 min at 56°C and subsequently alkylated with iodoacetamide (IAA) for 30 min at room temperature in the dark (final concentration 40 mM). Remaining IAA was quenched by DTT (final concentration 5 mM), and then Sequencing Grade Modified Trypsin (Promega) was added at the proportion 1:50 trypsin:protein ratio (w/w) following overnight incubation at 37°C. The samples were acidified adding trifluoroacetic acid (TFA) to a final concentration of 1% and digested peptides were desalted using C18 microcolumns (Thermo Fischer Scientific). Co-IP digested peptides were analyzed by Nano-HPLC-ESI-QUAD-TOF using a maXis 3G Bruker Daltonics (Central Analítica – IQ – USP). In brief, peptides were separated at a flow of 200 nl/min on a nanoAcquity UPLC® 1.8µm HSS T3 (75 µm x 200 mm) by reversed-phase chromatography, which was operated on a NanoAcquity Waters (Waters). The mobile phase was water/0.1% Formic Acid (A) and ACN/0.1% Formic Acid (B) during 240 minutes. The gradient was 2-30% phase B for 210 minutes, 30-85% B for 15 minutes, and 85-2% B for 15 minutes. The NanoAcquity Waters was coupled into a MAXIS 3G mass spectrometer (Bruker Daltonics) operating in positive ion mode. The mass spectrometer acquired a full MS scan at 60,000 full width half maximum (FWHM) resolution with a 350-2200 Da mass range. Top 10 most intense ions were selected from MS for Collision Induced Dissociation (CID) fragmentation (normalized collision energy: 6V). Exclusion time 120 seconds and 2 to 5 change range.

### MS/MS counting and label-free quantification

Bruker Q-TOF files were imported to MaxQuant version 1.6.17.0 for protein identification and quantification. For protein identification in MaxQuant, the database search engine Andromeda was used against Uniprot *Homo sapiens* (20,370 entries release). The following parameters were used: carbamidomethylation of cysteine (57.021464 Da) as a fixed modification, oxidation of methionine (15.994915 Da) and N-terminal acetylation protein (42.010565 Da) were selected as variable modifications. Enzyme specificity was set to full trypsin with maximum of two missed cleavages. The minimum peptide length was set to 7 amino acids. For label-free quantification, it was used “match between runs” feature in MaxQuant, which is able to identify the transfer between the samples based on the retention time and accurate mass, with a 0.7-minute match time window and 20 minutes’ alignment time window. Normalized MS/MS count was used as main parameter to further analyses. The mass spectrometry proteomics data is available at the ProteomeXchange Consortium (https://www.ebi.ac.uk/pride) via the PRIDE [58] partner repository with the dataset identifier PXD038402.

### Protein network and enrichment analysis

After the identification of peptides by normalized MS/MS count, proteins were selected as potential M2-2 binding partners by the following steps: (1) selection of enriched peptides (Fold Change > 1.5) or unique proteins for each independent experiment, excluding proteins present in negative controls, (2) selection of proteins identified in at least two out of three independent experiments. From this filtering, 72 proteins were selected as potential M2-2 interactors. These proteins were analyzed on STRING, for the generation of a functional protein-protein association network. Linking between network nodes was set for confidence, with minimum interaction score of 0.900. Analysis of enriched GO terms for Biological Process was performed on BiNGO (Cytoscape 3.9.1.), looking for overrepresented terms. Hypergeometric test was used for statistical analysis, with Benjamini-Hochberg False Discovery Rate correction, and significance set as 0.01.

### Western Blot

For protein analysis by western blot, cells were lysed in RIPA buffer (50 mM Tris-HCl pH 8.0, 150 mM NaCl, 1% Triton X-100, 0.5% sodium deoxycholate, 0.1% SDS) and quantified using the QuantiPro BCA Assay Kit (Sigma). Samples were then boiled in Laemmli Buffer for 10 min and submitted to SDS-PAGE, followed by transfer to 0.2 µm nitrocellulose membrane (Sigma) and blocking with 5% BSA (w/v) or 5% non-fat milk diluted in PBS-T (PBS 0.1% Tween 20). Primary and secondary antibodies were incubated overnight at 4°C and 1h at room temperature, respectively. After each incubation, membranes were rinsed three times for 5 min with PBS-T. HRP detection was performed with Clarity Western ECL Substrate (Bio-Rad). For protein normalization, membranes were stripped for 30 min at room temperature with stripping buffer (0.2 M glycine, 0.1% SDS, 1% Tween 20, pH 2.2) and washed three times with PBS-T before the next probing.

### Immunofluorescence staining

For immunofluorescence assays, cells were grown in glass coverslips at 70% confluence, and then submitted to transfection, infection, and treatments. 24h post treatments, cells were fixed with PBS containing 4% paraformaldehyde (w/v) for 15 min at 4°C, permeabilized with PBS 0.2% Triton X-100 for 15 min at 37°C and blocked with blocking solution (PBS, 0.1% Tween 20, 5% BSA) for 1h at 37°C. Primary and secondary antibodies were incubated in blocking solution overnight at 4°C and for 1h at 37°C, respectively, washing cells with PBS-T for three times between each step. Nuclei were stained with 4’,6-diamidino-2-phenylindole (DAPI) with secondary antibodies. Coverslips were mounted with Prolong Glass (Thermo Fisher Scientific), following curation for 24h in the dark at room temperature before imaging cells. For anti-calreticulin and anti-PABP staining, cells were fixed with ice-cold methanol for 10 min, following the same steps. Widefield microscopy was performed with a ZEISS Axio Vert.A1 microscope utilizing an objective Plan-NEOFLUAR 63x/NA 1.25 in oil immersion, and acquired images were analyzed on ImageJ (FIJI).

### Confocal microscopy

Confocal images were taken with a Zeiss LSM 780 NLO confocal microscope (Core Facility for Scientific Research – CEFAP-USP). Images were acquired with an objective α Plan-Apochromat 100x/NA 1.46 in oil immersion, using the following lasers: HeNe (543 nm), Argon (488 nm) and Diode (405 nm). Fluorescent signal was detected with a 32 channel GaAsP QUASAR detector with the following parameters: Alexa Fluor 594 (578 – 692 nm), Alexa Fluor 488 (491 – 587 nm), DAPI (412 – 491 nm). Pinhole was set to 1 airy unit, and z-stacks were taken with intervals of 290 nm. For deconvolution, theoretical PSF was generated, and images were processed using the DeconvolutionLab2 plugin [59] utilizing the Richardson-Lucy Total variation algorithm [60]. For all confocal immunofluorescences presented in this work, the images show a single z-stack. Images were contrast enhanced between the same experiments for better visualization.

### Reporter Luciferase Assay

HEK293T cells were co-transfected with pLPL (carrying Renilla and Firefly luciferases) and FLAG-EGFP, FLAG-M2-2 or EGFP-M2-2. After 24h, cells were rinsed and lysed with passive lysis buffer (Promega). Luminescence assay was performed with the Dual-Glo Luciferase Assay System (Promega) following manufacture’s recommendations, and luciferase activity was measured on GloMax Discover (Promega). For the reporter assay in infected cells, HEK293T cells were infected for 1h, and then transfected for 4h. Lysis was performed at 24 hpi.

### Statistical analysis

Statistics was performed on GraphPad Prism 9.4.0, and details of statistical tests used, error bars and p-values are described in figure legends. Statistical tests were only performed in data obtained in independent biological replicates, represented on graphs by colored dots, where each color indicates paired values.

## Acknowledgements

We are grateful to Cinthia L. Araújo for her previous work subcloning M2-1 on pcDNA3.1-FLAG. We acknowledge Mário C. Cruz for assistance with confocal microscopy and image analysis. We thank Marcus V. C. Baldo for discussions concerning statistical analysis. We are grateful to Carlos F. M. Menck for making his laboratory and reagents available for this work.

## Supporting information

**S1 Fig.**
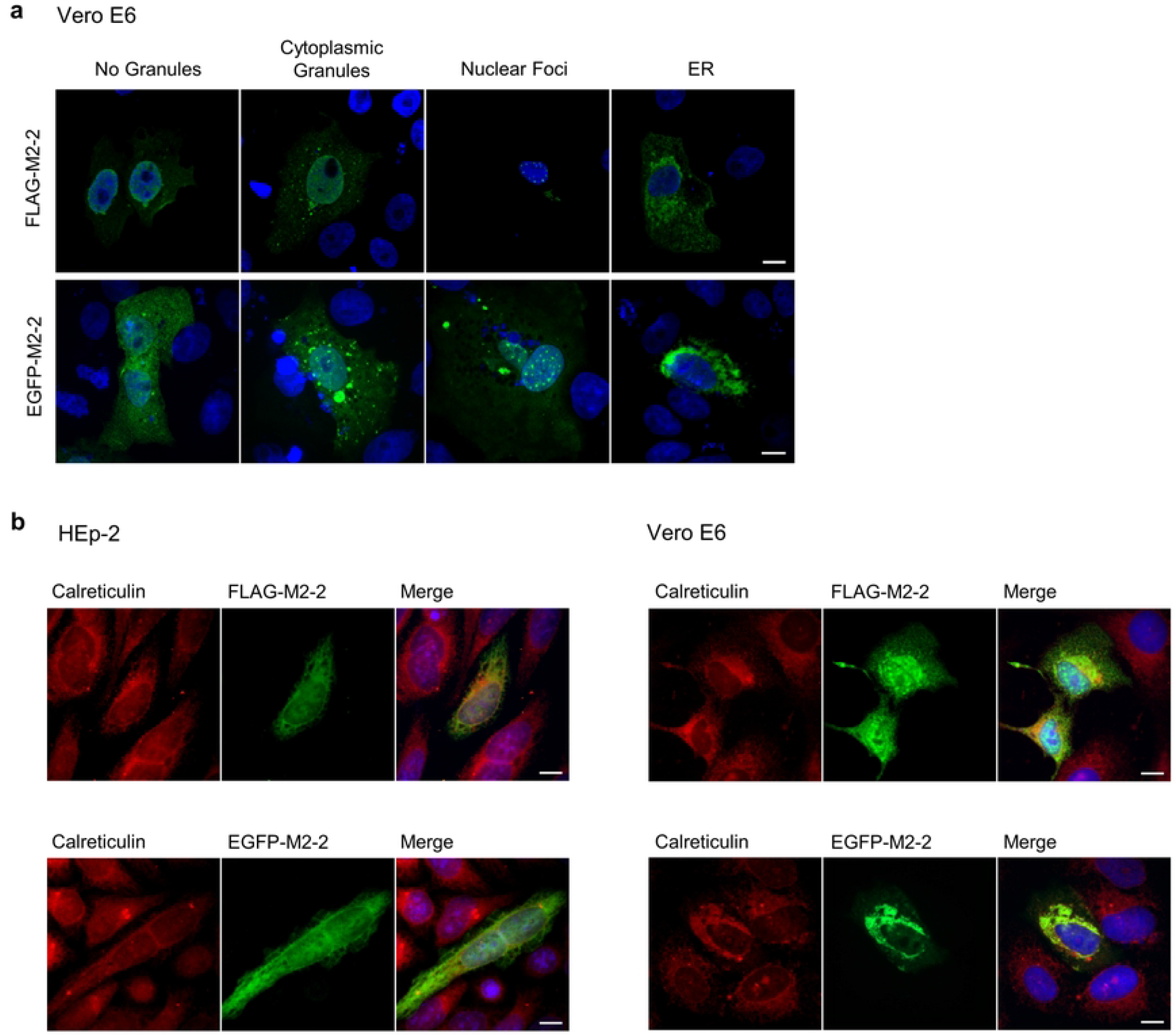
M2-2 colocalizes with the ER marker calreticulin in both HEp-2 and Vero E6 cells. (a) Immunofluorescence of FLAG-M2-2 and EGFP-M2-2 expressed for 24h in Vero E6 cells, presenting a distribution pattern similar to that observed in HEp-2 cells (Fig 1b). (b) Co-staining of M2-2 and the ER marker calreticulin, indicating that M2-2 could be recruited to these structures in different cell lines. All images were taken with a ZEISS Axio Vert.A1 microscope and are representative of three independent experiments. Scale bars 10 μm.

**S2 Fig.**
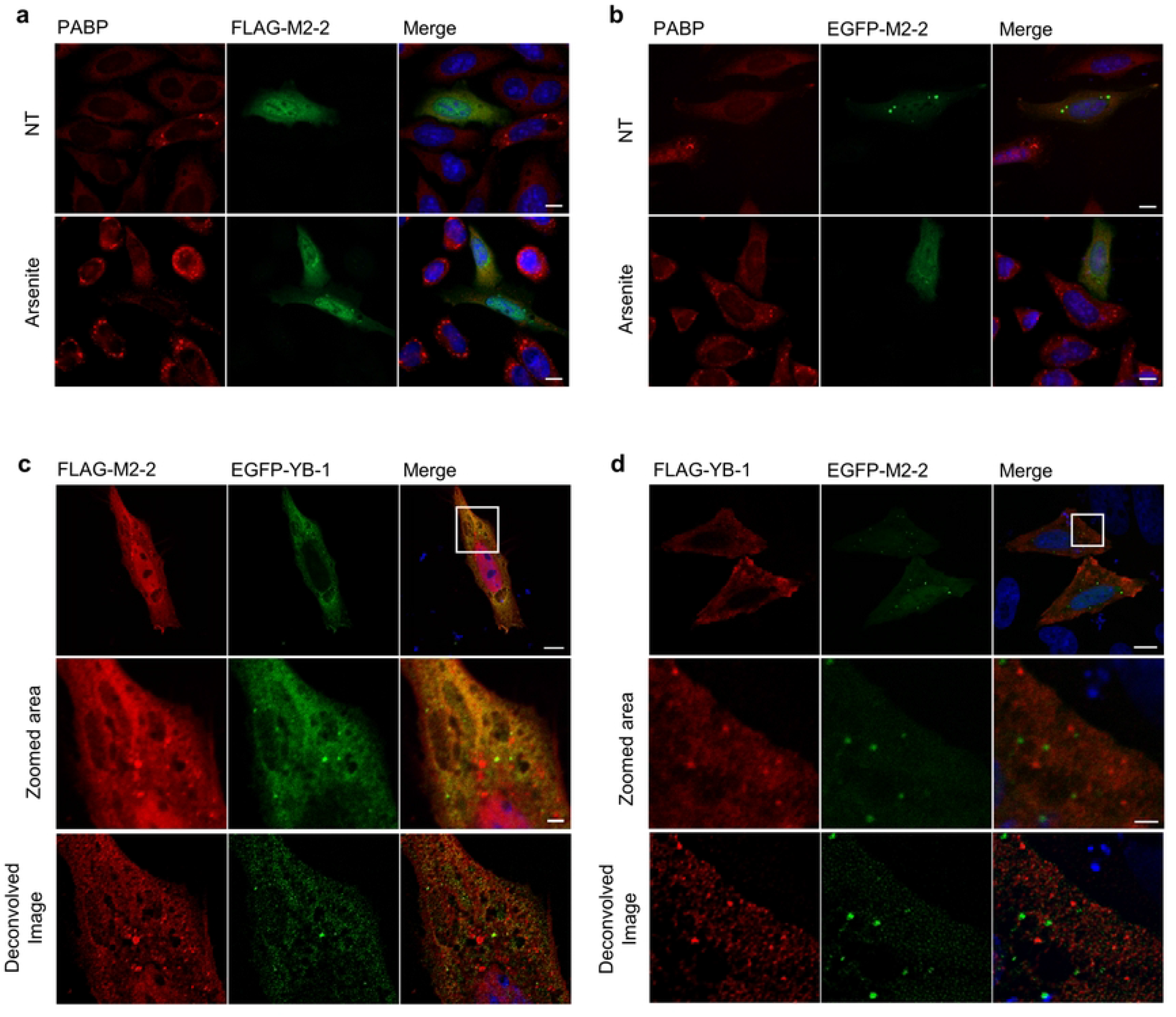
M2-2 does not colocalize with PABP or YB-1, markers of SGs and p-bodies. Widefield immunofluorescence of FLAG-M2-2 (a) and EGFP-M2-2 (b) expression in HEp-2 cells treated or not with arsenite (0.5 mM for 30 min) and stained against PABP, showing inhibition of SGs assembly. (c) Confocal immunofluorescence showing the co-expression of FLAG-M2-2 and EGFP-YB-1, marker of SGs and p-bodies. Zoomed area presents raw and deconvolved images, where M2-2 granules and p-bodies stained by EGFP-YB-1 can be distinguished as different structures. (d) Confocal immunofluorescence of cells co-expressing FLAG-YB-1 and EGFP-M2-2, as described in (c). All images are representative of three independent experiments. Scale bars are 10 µm for full size panels and 2 µm for zoomed areas.

**S3 Fig.**
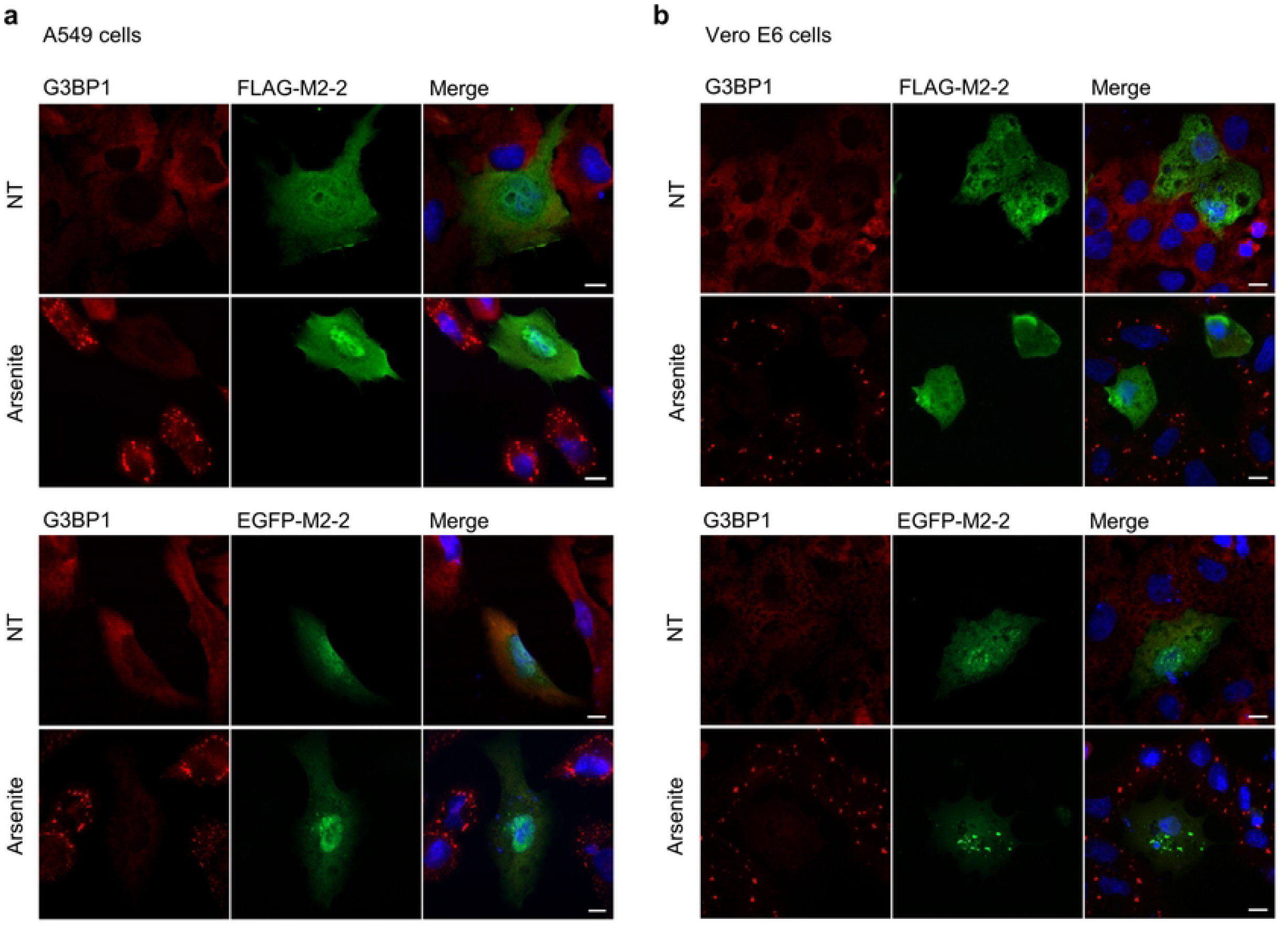
Inhibition of SGs assembly by M2-2 expression in different cell lines. Inhibition of SGs assembly, stained by G3BP1, is also shown in A549 (a) and Vero E6 cells (b), which is reproduced for both FLAG-M2-2 and EGFP-M2-2. Both proteins were expressed for 24h. Arsenite-treated cells (A549 – 0.5 mM for 30 min, Vero E6 – 1 mM for 1h) are indicated on the left. All images were taken with a ZEISS Axio Vert.A1 microscope and are representative of three independent experiments. Scale bars 10 µm.

**S4 Fig.**
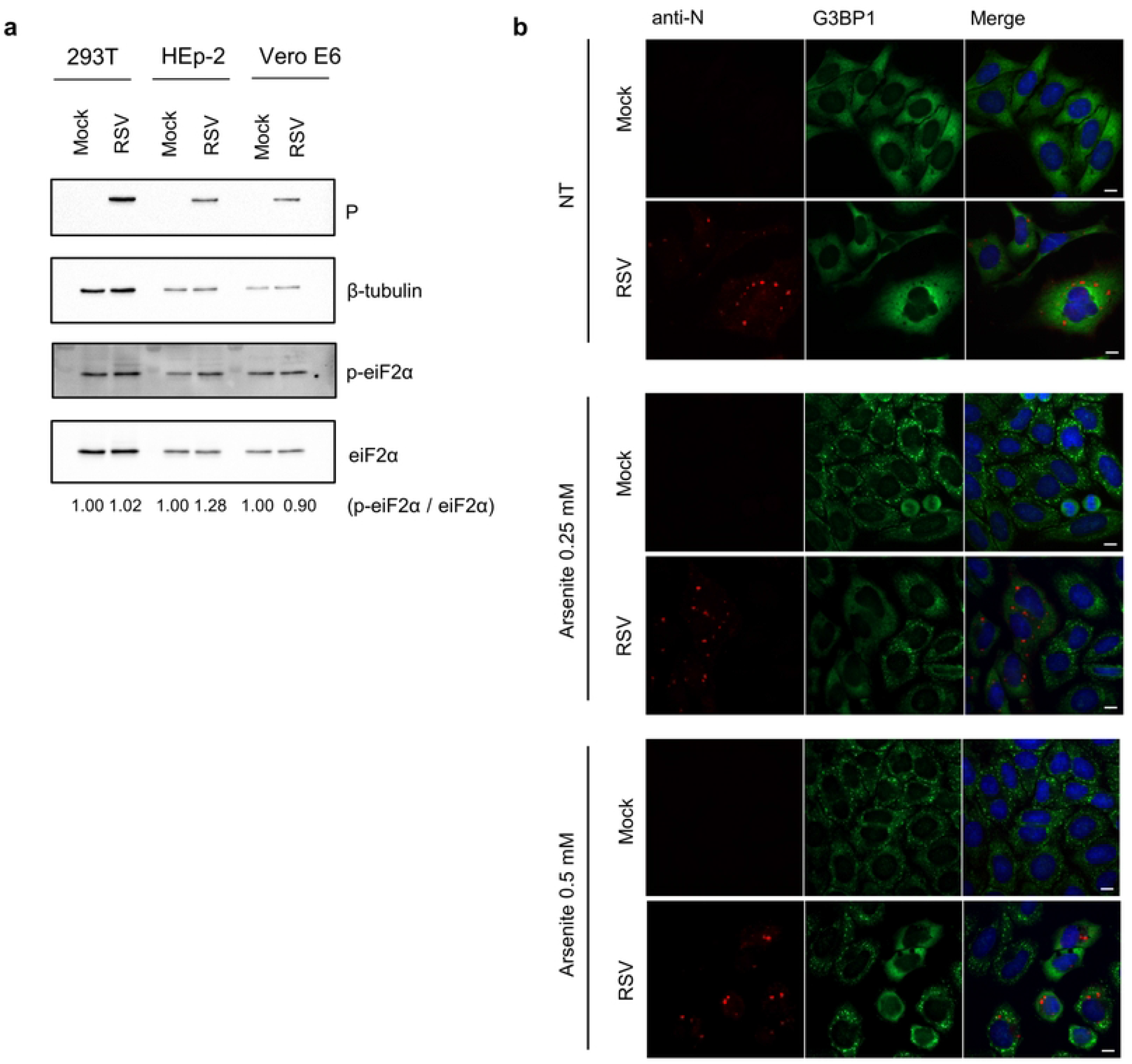
RSV infection does not trigger eiF2α phosphorylation or SGs assembly. (a) Western blot detection of phospho-eiF2α in HEK293T, HEp-2 and Vero E6 cells mock, or RSV infected for 24h. Phospho-eiF2α / total eiF2α ratios are shown below. Infection is indicated by detection of the RSV P protein. (b) HEp-2 cells were mock, or RSV infected for 24h and submitted or not to arsenite treatment for 30 min at the indicated concentrations. SGs were stained by G3BP1 while RSV infected cells were detected with anti-N. Impairment of SGs assembly in infected cells can be seen in all panels. Images were taken with a widefield ZEISS Axio Vert.A1 microscope and are representative of three independent experiments. Scale bars 10 μm.

**S5 Fig.**
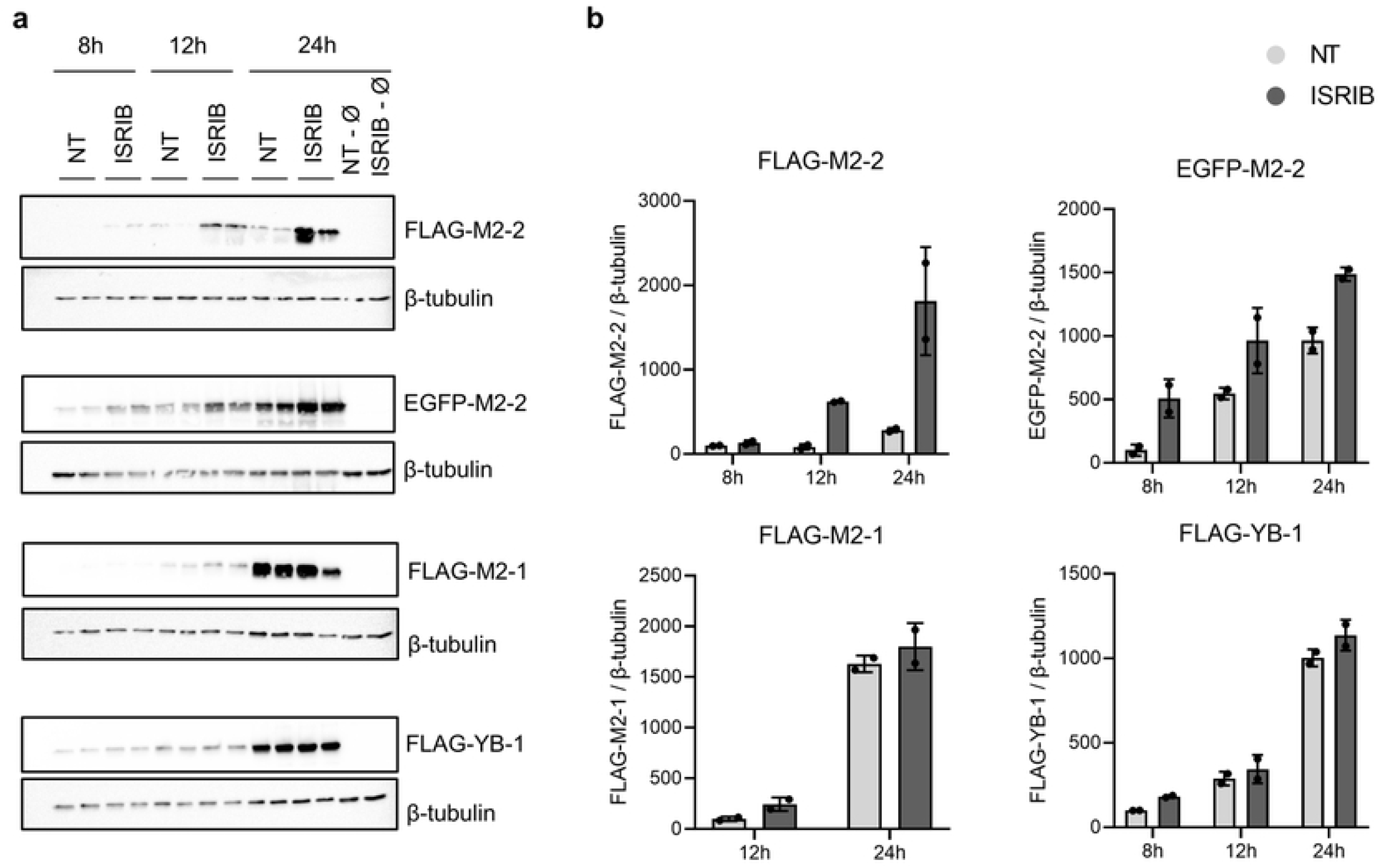
ISRIB treatment enhances M2-2 expression. (a) Time-course expression of FLAG-M2-2 and EGFP-M2-2 in HEK293T performed in duplicate. Cells were ISRIB treated (200 nM) or not (NT – non-treated) and collected at indicated times for western blot analysis. As controls, cells were transfected with FLAG-M2-1 or FLAG-YB-1. Quantification of detected bands is shown in graphs on (b), showing the specific effect of ISRIB in the expression of M2-2.

**S6 Fig.**
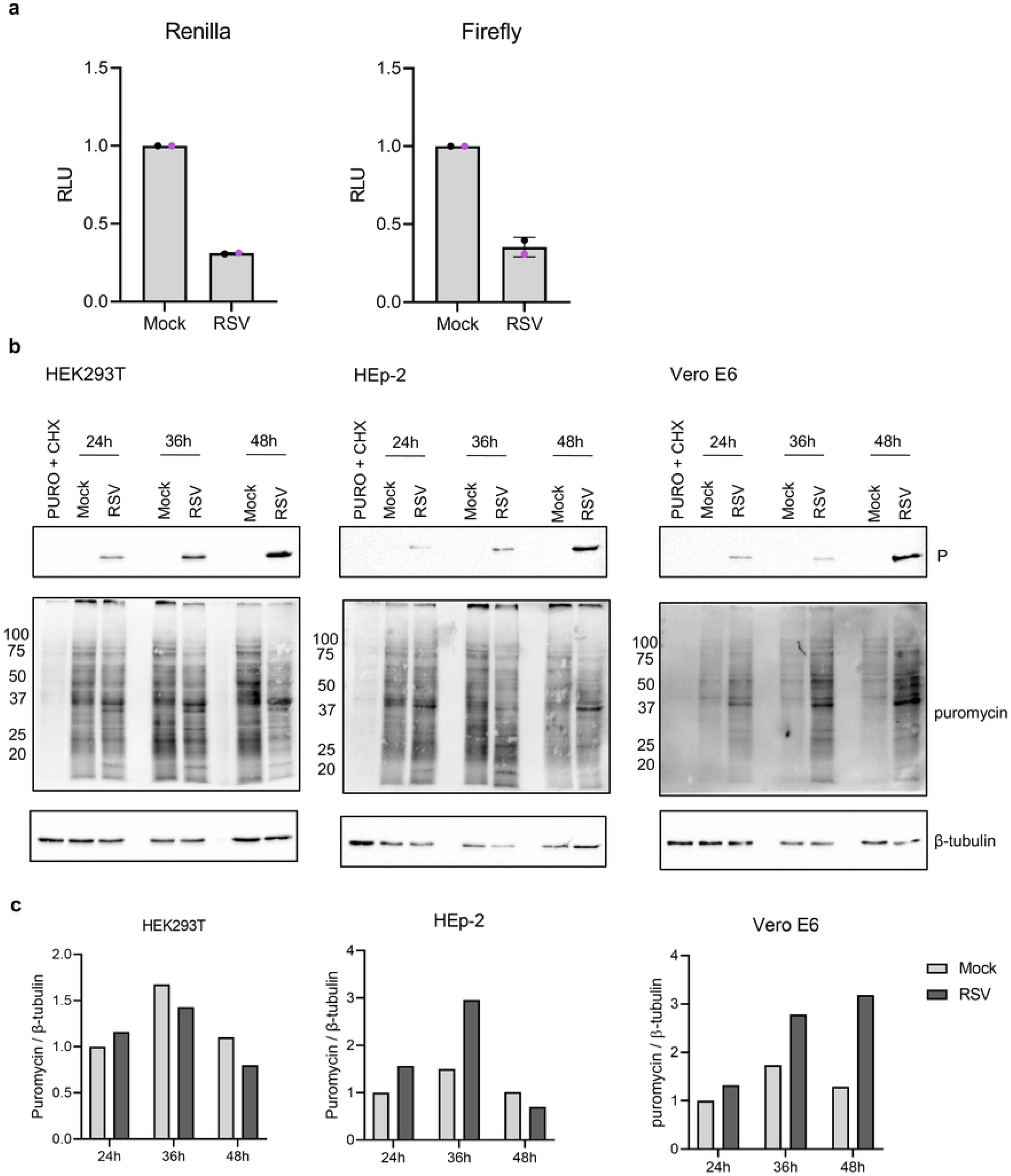
RSV infection inhibits 5’cap and IRES-dependent translation initiations but does not reduce the global translation rate. (a) HEK293T cells were mock or RSV infected, and 1 hpi cells were transfected with the vector for expression of reporter luciferases. 24 hpi, cells were lysed, and luminescence activity was evaluated, showing inhibition of both Renilla (5’ cap) and Firefly (IRES) luciferases. Colored dots in the graphs indicate individual means of two independent experiment performed in triplicate (n=2). Error bars show standard deviation. (b) SUnSET assay performed in HEK293T, HEp-2 and Vero E6 cells. Cells were mock or RSV infected and treated with puromycin (10 µg/mL) at indicated times. As negative control, cells were treated with puromycin and cycloheximide (100 µg/mL). Molecular weight of puromycin bands is shown on the left, and labeled proteins are indicated on the right. Graphs on (c) show quantified puromycin detection normalized to β-tubulin.

**S7 Fig.**
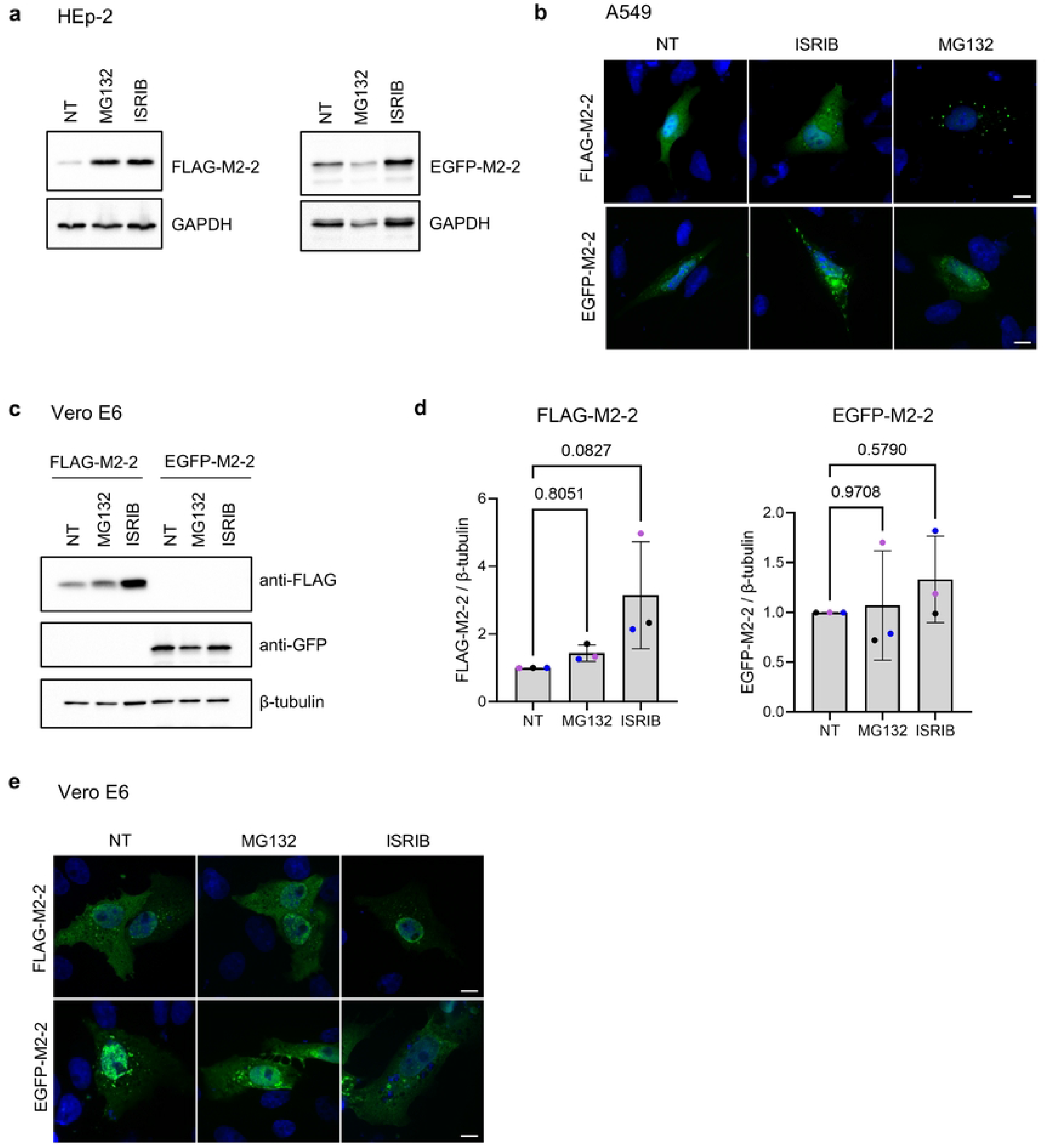
MG132 effects on M2-2 expression in HEp-2, A549, and Vero E6 cells. (a) Expression of FLAG and EGFP-M2-2 in HEp-2 cells treated or not with ISRIB (200 nM) and MG132 (5 µM) for 20h. (b) Cellular distribution of FLAG and EGFP-M2-2 in A549 cells treated or not with ISRIB (200 nM for 20h) and MG132 (5 µM for 2h), showing increase in the number of FLAG-M2-2 granules under MG132 treatment. (c) FLAG-M2-2 and EGFP-M2-2 expression in Vero E6 cells for 24h. Cells were treated with ISRIB or MG132, as in (a). Normalized expression of the proteins in (c) is shown in (d), where each colored dot represents values from the same independent experiment (n=3). Differences between means were accessed by one-way anova followed by Dunnett’s multiple comparisons test. Error bars indicate standard deviation, and p-values are shown on the graph. (e) Cellular distribution of FLAG-M2-2 and EGFP-M2-2 in Vero E6 cells, showing no major changes between indicated treatments. All immunofluorescence images were taken with a widefield ZEISS Axio Vert.A1 microscope and are representative of three independent experiments. Scale bars 10 μm.

**S8 Fig.**
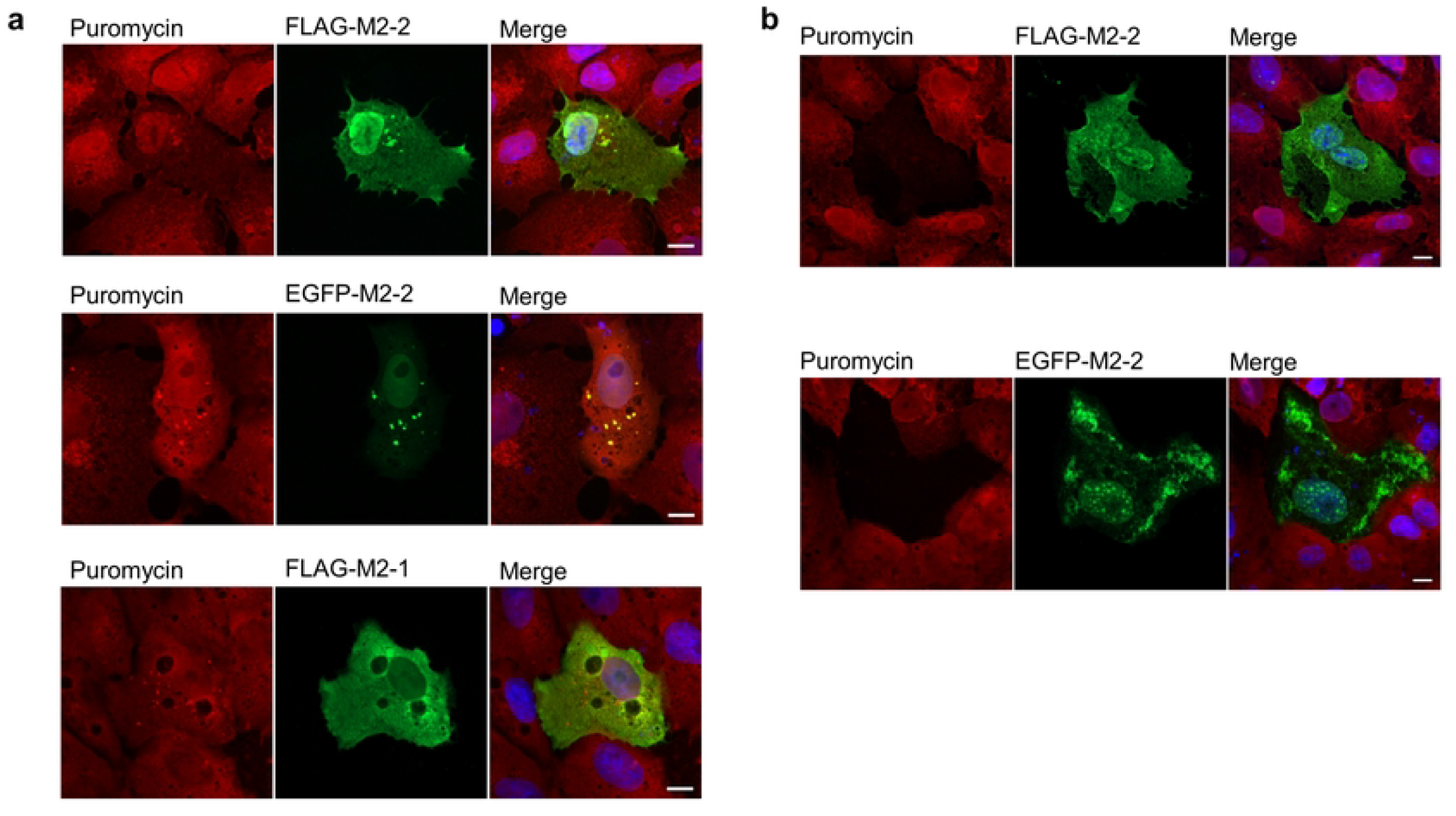
M2-2 colocalizes with DRiPs and inhibits translation in Vero E6 cells. (a) FLAG-M2-2, EGFP-M2-2 and FLAG-M2-1 were expressed in Vero E6 cells and treated with puromycin (10 µg/mL) and MG132 (5 µM) for 2h. Both FLAG and EGFP-M2-2 colocalized with puromycin granules in these cells, with no recruitment of FLAG-M2-1 to puromycin granules. (b) Images show cells were M2-2 expression could inhibit puromycin incorporation, indicating that M2-2 ability to hamper translation is kept in this cell line. All images were taken with a ZEISS Axio Vert.A1 microscope and are representative of two independent experiments. Scale bars 10 μm.

**S9 Fig.**
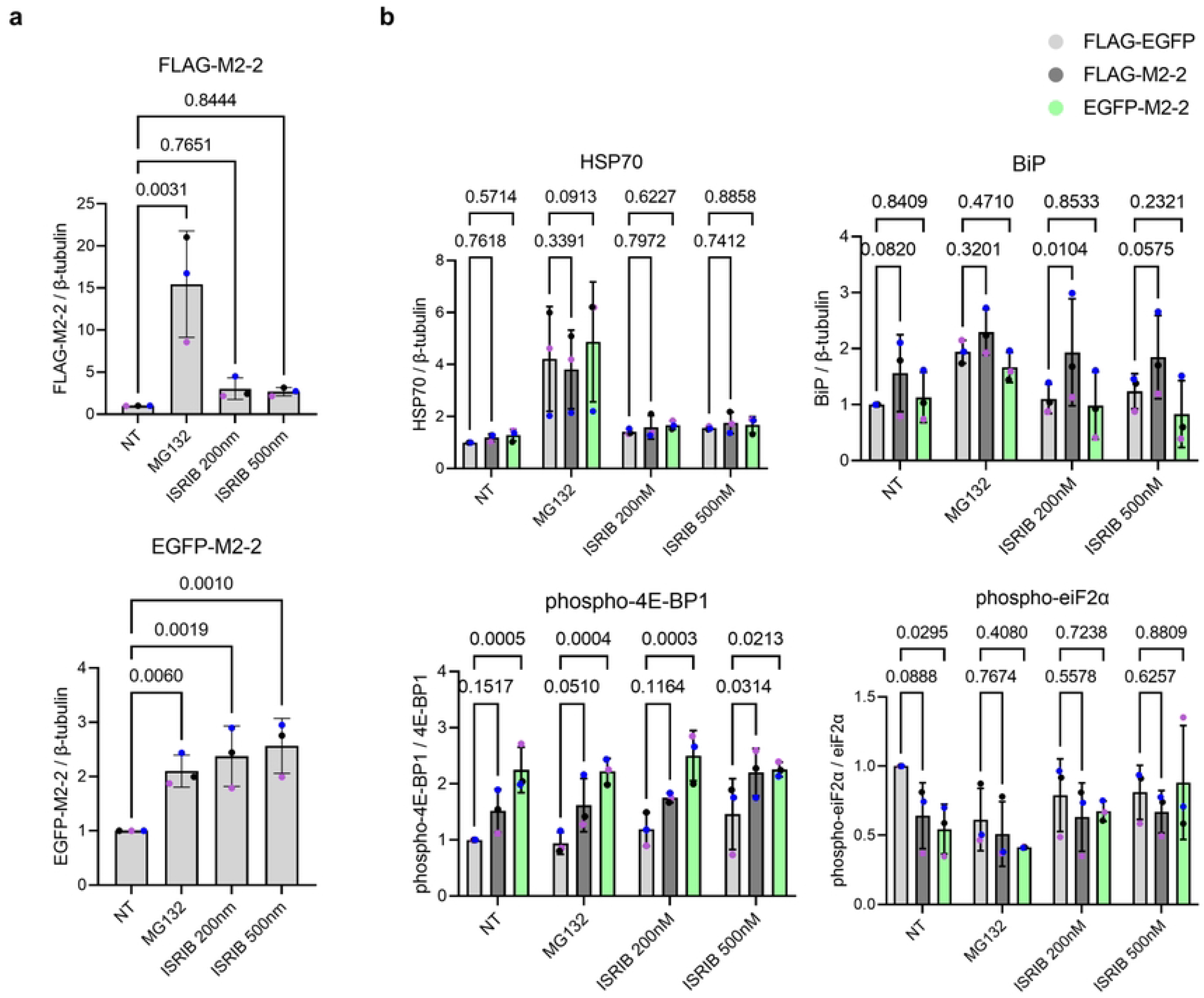
M2-2 does not enhance the expression of chaperones but modulates the phosphorylation of translation effectors. (a) Quantitative analysis of FLAG-M2-2 and EGFP-M2-2 expression under MG132 and ISRIB treatments shown in Figure 7d. (b) Quantitative analysis of the expression of proteostatic stress markers (HSP70 and BiP), and modulators of translation (phospho-4E-BP1 and phospho-eiF2α), from western blots shown in Figure 7d. Statistical differences were evaluated by one-way anova (a) or two-way anova (b), both followed by Dunnett’s multiple comparisons test. Colored dots in all graphs indicate individual values from the same independent experiment (n=3). Error bars indicate standard deviation and p-values are shown in the graphs.

**S10 Fig.**
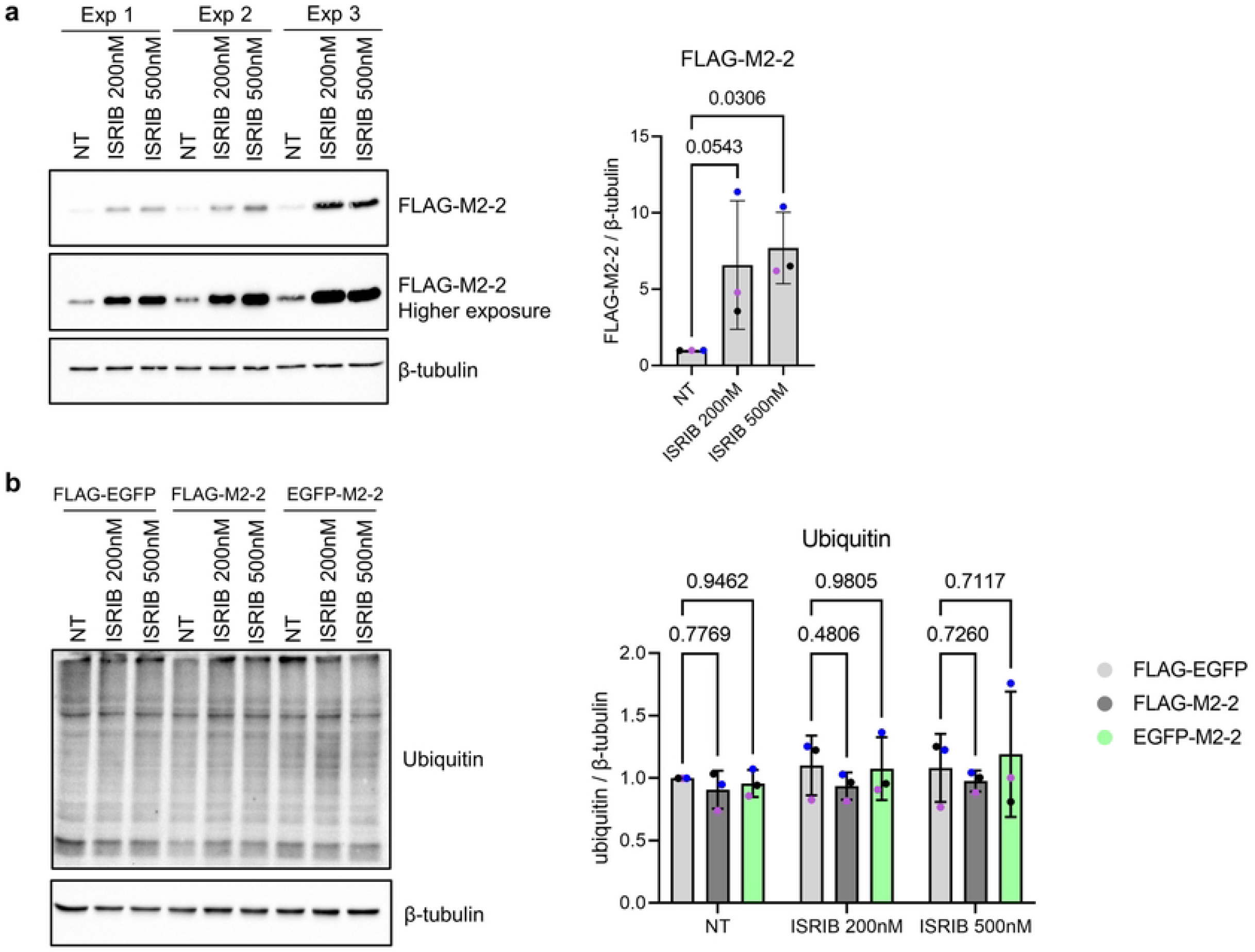
Evaluation of FLAG-M2-2 and ubiquitin detection under ISRIB treatment. (a) To better evaluate FLAG-M2-2 expression in the absence of saturated bands, samples of three independent experiments from NT and ISRIB (200 nM and 500 nM) treatments were analyzed by western blot (same samples from Fig 7d). Graph of quantified bands is shown on the right. (b) Comparison of ubiquitin levels between non-treated cells and ISRIB treatments in the absence of MG132 treated samples. Quantitative analysis of three independent experiments is shown on the graph (right). Statistical differences were evaluated by one-way anova (a) or two-way anova (b), both followed by Dunnett’s multiple comparisons test. Colored dots in graphs indicate individual values from paired independent experiments (n=3). Error bars indicate standard deviation and p-values are shown in the graphs.

**S1 Table. Mass spectrometry and enrichment analysis**.

**S1 File. DNA sequence of optimized genes**.

## References

1. Afonso CL, Amarasinghe GK, Bányai K, Bào Y, Basler CF, Bavari S, et al. Taxonomy of the order Mononegavirales: update 2016. Archives of Virology. 2016; 161, 2351–2360.

2. Collins PL, Melero JA. Progress in understanding and controlling respiratory syncytial virus: still crazy after all these years. Virus Research. 2011; 162, 80–99.

3. Bohmwald K, Espinoza JA, Rey-Jurado E, Gómez RS, González PA, Bueno SM, et al. Human Respiratory Syncytial Virus: infection and pathology. Semin Respir Crit Care Med. 2016; 37, 522–37.

4. Bakker SE, Duquerroy S, Galloux M, Loney C, Conner E, Eléouët JF, et al. The respiratory syncytial virus nucleoprotein-RNA complex forms a left-handed helical nucleocapsid. J Gen Virol. 2013; 94, 1734–1738.

5. Galloux M, Tarus B, Blazevic I, Fix J, Duquerroy S, Eléouët JF. Characterization of a viral phosphoprotein binding site on the surface of the respiratory syncytial nucleoprotein. J Virol. 2012; 86, 8375–87.

6. Sourimant J, Rameix-Welti MA, Gaillard AL, Chevret D, Galloux M, Gault E, et al. Fine mapping and characterization of the L-polymerase-binding domain of the respiratory syncytial virus phosphoprotein. J Virol. 2015; 89, 4421–33.

7. Bermingham A, Collins PL. The M2-2 protein of human respiratory syncytial virus is a regulatory factor involved in the balance between RNA replication and transcription. Proc Natl Acad Sci U S A. 1999; 96, 11259–64.

8. Fearns R, Collins PL. Role of the M2-1 transcription antitermination protein of respiratory syncytial virus in sequential transcription. J Virol. 73, 5852–64 (1999).

9. Cowton VM, McGivern DR, Fearns R. Unravelling the complexities of respiratory syncytial virus RNA synthesis. J Gen Virol. 2006; 87, 1805–21.

10. Pereira N, Cardone C, Lassoued S, Galloux M, Fix J, Assrir N, et al. New insights into structural disorder in human respiratory syncytial virus phosphoprotein and implications for binding of protein partners. J Biol Chem. 2017; 292, 2120–2131.

11. Galloux M, Risso-Ballester J, Richard CA, Fix J, Rameix-Welti MA, Eléouët JF. Minimal elements required for the formation of Respiratory Syncytial Virus cytoplasmic inclusion bodies in vivo and in vitro. mBio. 2020; 11, e01202–20.

12. Rincheval V, Lelek M, Gault E, Bouillier C, Sitterlin D, Blouquit-Laye S, et al. Functional organization of cytoplasmic inclusion bodies in cells infected by respiratory syncytial virus. Nature Communications. 2017; 8, 563.

13. Richard CA, Rincheval V, Lassoued S, Fix J, Cardone C, Esneau C, et al. RSV hijacks cellular protein phosphatase 1 to regulate M2-1 phosphorylation and viral transcription. PLoS Pathog. 2018; 14, e1006920.

14. Jin H, Cheng X, Zhou HZ, Li S, Seddiqui A. Respiratory syncytial virus that lacks open reading frame 2 of the M2 gene (M2-2) has altered growth characteristics and is attenuated in rodents. J Virol. 2000; 74, 74–82.

15. Blanchard EL, Braun MR, Lifland AW, Ludeke B, Noton SL, Vanover D, et al. Polymerase-tagged respiratory syncytial virus reveals a dynamic rearrangement of the ribonucleocapsid complex during infection. PLoS Pathog. 2020; 16, e1008987.

16. Goswami R, Majumdar T, Dhar J, Chattopadhyay S, Bandyopadhyay SK, Verbovetskaya V, et al. Viral degradasome hijacks mitochondria to suppress innate immunity. Cell Res. 2013; 23, 1025–42.

17. Pei J, Beri NR, Zou AJ, Hubel P, Dorando HK, Bergant V, et al. Nuclear-localized human respiratory syncytial virus NS1 protein modulates host gene transcription. Cell Rep. 2021; 37, 109803.

18. Li HM, Ghildyal R, Hu M, Tran KC, Starrs LM, Mills J, et al. Respiratory Syncytial Virus matrix protein-chromatin association is key to transcriptional inhibition in infected cells. Cells. 2021; 10, 2786.

19. Lifland AW, Jung J, Alonas E, Zurla C, Crowe JE Jr, Santangelo PJ. Human respiratory syncytial virus nucleoprotein and inclusion bodies antagonize the innate immune response mediated by MDA5 and MAVS. J Virol. 2012; 86, 8245–58.

20. Munday DC, Wu W, Smith N, Fix J, Noton SL, Galloux M, et al. Interactome analysis of the human respiratory syncytial virus RNA polymerase complex identifies protein chaperones as important cofactors that promote L-protein stability and RNA synthesis. J Virol. 2015; 89, 917–30.

21. Bouillier C, Cosentino G, Léger T, Rincheval V, Richard CA, Desquesnes A, et al. The interactome analysis of the Respiratory Syncytial Virus protein M2-1 suggests a new role in viral mRNA metabolism post-transcription. 2019; Sci Rep. 9, 15258.

22. Cervantez-Ortiz SL, Cuervo NZ, Grandvaux N. Respiratory Syncytial Virus and cellular stress responses: impact on replication and physiopathology. Viruses. 2016; 8, 124.

23. Zhang Q, Sharma NR, Zheng ZM, Chen M. Viral regulation of RNA granules in infected cells. Virol Sin. 2019; 34, 175–191.

24. Jain S, Wheeler JR, Walters RW, Agrawal A, Barsic A, Parker R. ATPase-modulated stress granules contain a diverse proteome and substructure. Cell. 2016; 164, 487–98.

25. Sidrauski C, McGeachy AM, Ingolia NT, Walter P. The small molecule ISRIB reverses the effects of eIF2α phosphorylation on translation and stress granule assembly. eLife. 2015; 4, e05033.

26. Zyryanova AF, Weis F, Faille A, Alard AA, Crespillo-Casado A, Sekine Y, et al. Binding of ISRIB reveals a regulatory site in the nucleotide exchange factor eIF2B. Science. 2018; 359, 1533–1536.

27. Yang WH, Bloch DB. Probing the mRNA processing body using protein macroarrays and “autoantigenomics”. RNA. 2007; 13, 704–12.

28. tanaka T, Ohashi S, Kobayashi S. Roles of YB-1 under arsenite-induced stress: Translational activation of HSP70 mRNA and control of the number of stress granules. Biochim Biophys Acta. 2014; 1840, 985–92.

29. Groskreutz DJ, Babor EC, Monick MM, Varga SM, Hunninghake GW. Respiratory syncytial virus limits alpha subunit of eukaryotic translation initiation factor 2 (eIF2alpha) phosphorylation to maintain translation and viral replication. J Biol Chem. 2010; 285, 24023–31.

30. Hanley LL, McGivern DR, Teng MN, Djang R, Collins PL, Fearns R. Roles of the respiratory syncytial virus trailer region: effects of mutations on genome production and stress granule formation. Virology. 2010; 406, 241–52.

31. Lindquist ME, Lifland AW, Utley TJ, Santangelo PJ, Crowe JE Jr. Respiratory syncytial virus induces host RNA stress granules to facilitate viral replication. J Virol. 2010; 84, 12274–84.

32. Rabouw HH, Langereis MA, Anand AA, Viser LJ, Groot RJ, Walter P, et al. Small molecule ISRIB suppresses the integrated stress response within a defined window of activation. Proc Natl Acad Sci U S A. 2019; 116, 2097–2102.

33. Schmidt EK, Clavarino G, Ceppi M, Pierre P. SUnSET, a nonradioactive method to monitor protein synthesis. Nat Methods. 2009; 6, 275–7.

34. Gerlitz G, Jagus R, Elroy-Stein O. Phosphorylation of initiation factor-2 alpha is required for activation of internal translation initiation during cell differentiation. Eur J Biochem. 2002; 269, 2810–9.

35. Gallo S, Ricciardi S, Manfrini N, Pesce E, Oliveto S, Calamita P, et al. RACK1 specifically regulates translation through its binding to ribosomes. Mol Cell Biol. 2018; 38, e00230–18.

36. LaFontaine E, Miller CM, Permaul N, Martin ET, Fuchs G. Ribosomal protein RACK1 enhances translation of poliovirus and other viral IRESs. Virology. 2020; 545, 53–62.

37. Jackson RJ, Hellen CUT, Pestova TV. The mechanism of eukaryotic translation initiation and principles of its regulation. Nat Rev Mol Cell Biol. 2010; 11, 113–27.

38. Lee ASY, Burdeinick-Kerr R, Whelan SPJ. A ribosome-specialized translation initiation pathway is required for cap-dependent translation of vesicular stomatitis virus mRNAs. Proc Natl Acad Sci U S A. 2013; 110, 324–9.

39. Cockman E, Anderson P, Ivanov P. TOP mRNPs: molecular mechanisms and principles of regulation. Biomolecules. 2020; 10, 969.

40. Banerjee AK, Blanco MR, Bruce EA, Honson DD, Chen LM, Chow A, et al. SARS-CoV-2 disrupts splicing, translation, and protein trafficking to suppress host defenses. Cell. 2020; 183, 1325–1339.

41. Berglund P, Finzi D, Bennick JR, Yewdell JW. Viral alteration of cellular translational machinery increases defective ribosomal products. J Virol. 2007; 81, 7220–9.

42. DiGiuseppe S, Rollins MG, Bartom ET, Walsh D. ZNF598 plays distinct roles in interferon-stimulated gene expression and poxvirus protein synthesis. Cell Rep. 2018; 23, 1249–1258.

43. Sundaramoorthy E, Ryan AP, Fulzele A, Leonard M, Daugherty MD, Bennett EJ. Ribosome quality control activity potentiates vaccinia virus protein synthesis during infection. J Cell Sci. 2021; 134, jcs257188.

44. Latorre V, Geller R. Identification of cytoplasmic chaperone networks relevant for Respiratory Syncytial Virus replication. Front Microbiol. 2022; 13, 880394.

45. Duttler S, Pechmann S, Frydman J. Principles of cotranslational ubiquitination and quality control at the ribosome. Mol Cell. 2013; 50, 379–93.

46. Richter JD, Coller J. Pausing on polyribosomes: make way for elongation in translational control. Cell. 2015; 163, 292–300.

47. Sundaramoorthy E, Leonard M, Mak R, Liao J, Fulzele A, Bennett EJ. ZNF598 and RACK1 regulate mammalian ribosome-associated quality control function by mediating regulatory 40S ribosomal ubiquitylation. Mol Cell. 2017; 65, 751–760.

48. Lang WH, Calloni G, Vabulas RM. Polylysine is a proteostasis network-engaging structural determinant. J Proteome Res. 17, 1967–1977 (2018).

49. Joazeiro CAP. Mechanisms and functions of ribosome-associated protein quality control. Nat Rev Mol Cell Biol. 2019; 20, 368–383.

50. Liu XD, Ko S, Xu Y, Fattah EA, Xiang Q, Jagannath C, et al. Transient aggregation of ubiquitinated proteins is a cytosolic unfolded protein response to inflammation and endoplasmic reticulum stress. J Biol Chem. 2012; 287, 19687–98.

51. Mediani L, Guillén-Boixet J, Vinet J, Franzmann TM, Bigi I, Mateju D, et al. Defective ribosomal products challenge nuclear function by impairing nuclear condensate dynamics and immobilizing ubiquitin. EMBO J. 2019; 38, e101341.

52. Osada N, Kohara A, Yamaji T, Hirayama N, Kasai F, Sekizuka T, et al. The genome landscape of the African Green Monkey kidney-derived Vero cell line. DNA Res. 2014; 21, 673–683.

53. Biggiogera M, Cisterna B, Spedito A, Vecchio L, Malatesta M. Perichromatin fibrils as early markers of transcriptional alterations. Differentiation. 2008; 76, 57–65.

54. Galganski L, Urbanek MO, Krzyzosiak WJ. Nuclear speckles: molecular organization, biological function and role in disease. Nucleic Acids Res. 2017; 45, 10350–10368.

55. Yan LL, Zaher HS. Ribosome quality control antagonizes the activation of the integrated stress response on colliding ribosomes. Mol Cell. 2021; 81, 614–628.

56. Meyer C, Garzia A, Morozov P, Molina H, Tuschl T. The G3BP1-family-USP10 deubiquitinase complex rescues ubiquitinated 40S subunits of ribosomes stalled in translation from lysosomal degradation. Mol Cell. 2020; 77, 1193–1205.

57. Cho JY, Akbarali Y, Zerbini LF, Gu X, Boltax J, Wang Y, et al. Isoforms of the Ets transcription factor NERF/ELF-2 physically interact with AML1 and mediate opposing effects on AML1-mediated transcription of the B cell-specific blk gene. J Biol Chem. 2004; 279, 19512–22.

58. Perez-Riverol Y, Csordas A, Bai J, Bernal-Llinares M, Hewapathirana S, Kundu DJ, et al. The PRIDE database and related tools and resources in 2019: improving support for quantification data. Nucleic Acids Res. 2019; 47, D442–D450.

59. Sage D, Donati L, Soulez F, Fortun D, Schmit G, Seitz A, et al. DeconvolutionLab2: an open-source software for deconvolution microscopy. Methods. 2017; 115, 28–41.

60. Dey N, Blanc-Feraud L, Zimmer C, Roux P, Kam Z, Olivo-Marin JC, et al. Richardson-Lucy algorithm with total variation regularization for 3D confocal microscope deconvolution. Microsc Res Tech. 2006; 69, 260–6.

